# Single-cell profiling of bronchoalveolar cells reveals a Th17 signature in neutrophilic severe equine asthma

**DOI:** 10.1101/2023.07.04.547614

**Authors:** Sophie E. Sage, Tosso Leeb, Vidhya Jagannathan, Vinzenz Gerber

**Affiliations:** Swiss Institute of Equine Medicine, Department of Clinical Veterinary Medicine, Vetsuisse Faculty, University of Bern; Bern, Switzerland; Institute of Genetics, Vetsuisse Faculty, University of Bern; Bern, Switzerland; Clinical Diagnostic Laboratory, Department of Clinical Veterinary Medicine, Vetsuisse Faculty, University of Bern; Bern, Switzerland; Next Generation Sequencing Platform, University of Bern, Bern, Switzerland

## Abstract

Severe equine asthma (SEA) shares clinical and pathological features with human neutrophilic asthma, serving as a rare natural model for this condition. To uncover the elusive immune mechanisms driving SEA, we performed single-cell mRNA sequencing (scRNA-seq) on cryopreserved bronchoalveolar cells from 11 Warmblood horses, five controls and six with SEA. We identified six major cell types, showing significant heterogeneity and novel subtypes. Notably, we observed monocyte-lymphocyte complexes and detected a robust Th17 signature in SEA, with *CXCL13* upregulation in intermediate monocytes. Asthmatic horses exhibited expansion of the B cell population, Th17 polarization of the T cell populations, and dysregulation of genes associated with T cell function. Neutrophils demonstrated enhanced migratory capacity and heightened aptitude for neutrophil extracellular trap formation. These findings provide compelling evidence for a predominant Th17 immune response in neutrophilic SEA, driven by dysregulation of monocyte and T cell genes. The dysregulated genes identified through scRNA-seq have potential as biomarkers and therapeutic targets for SEA and provide insights into human neutrophilic asthma.

**One Sentence Summary:** Single-cell mRNA sequencing identifies a predominant Th17-mediated immune response in severe equine asthma

## INTRODUCTION

Equine asthma is a common respiratory disease of the horse characterized by bronchoconstriction, mucus production and bronchospasm (*1*). Its severe form, severe equine asthma (SEA), presents with increased breathing effort at rest, airway remodeling and in most cases, airway neutrophilia (*1*). Equine asthma is a very active field of research, in part because of its negative impact on animal welfare and the horse industry but also due to its similarities with human asthma, making it a rare natural animal model (*2–4*). In contrast to murine models with experimentally induced airway inflammation, horses develop asthma under natural conditions. Their longer lifespan enables the study of disease progression, particularly airway remodeling. Furthermore, their size facilitates the collection of lower airway samples. For instance, the collection of bronchoalveolar lavage fluid (BALF) is a routine procedure in horses, in contrast to humans and conventional laboratory animal models. Although promising asthma drugs have been identified based on murine studies, their limited clinical efficacy when applied to humans (*4*) may be attributed to disparities in the underlying pathophysiological mechanisms between experimentally induced and naturally occurring diseases.

In humans, asthma is considered an umbrella diagnosis encompassing a plethora of diseases with distinct pathophysiologic mechanisms (so-called endotypes). The advent of omics technologies has begun to unveil the diversity of human asthma endotypes (*5*, *6*). SEA shares clinical and pathological features with several human asthma endotypes, including allergic, non-allergic and late-onset asthma (*2*). Because they are exposed to high levels of dust in stables, horses represent an ideal model for organic dust-induced asthma of agricultural workers (*7*). While SEA has been mainly attributed to a Th2 response, there have also been reports of predominant Th1 and mixed Th1/Th2 phenotypes (*8*). Furthermore, the Th17 pathway, typically associated with autoimmune diseases, has been implicated (*9–12*). The complexity of the disease and limitations of experimental techniques may have contributed to these inconsistent findings. To address this knowledge gap, we leveraged the emerging single-cell mRNA sequencing (scRNA-seq) technology to dissect the immune mechanisms of SEA at the single-cell level.

In a previous experiment, we demonstrated that scRNA-seq can be successfully applied to fresh frozen equine BALF cells (*13*). Here, we employed the 10X Genomics droplet-based scRNA-seq technology to sequence BALF cells from six horses with SEA and five control horses. Our analysis revealed a distinct transcriptomic signature in several cell types from asthmatic horses, with monocytes, alveolar macrophages, and T cells exhibiting a clear Th17 polarization.

## RESULTS

### Single-cell landscape of bronchoalveolar lavage fluid from asthmatic and control horses

We analyzed the scRNA-seq data obtained from the BALF cells collected from six asthmatic and five control horses. Detailed characteristics of the study population can be found in Table 1. As anticipated, the Horse Owner Assessed Respiratory Signs Index (HOARSI) score and BALF neutrophil count were significantly different between the two groups (Table 1). Furthermore, the clinical score and tracheal mucus score exhibited significant differences, confirming appropriate phenotyping (*1*).

**Table 1:**
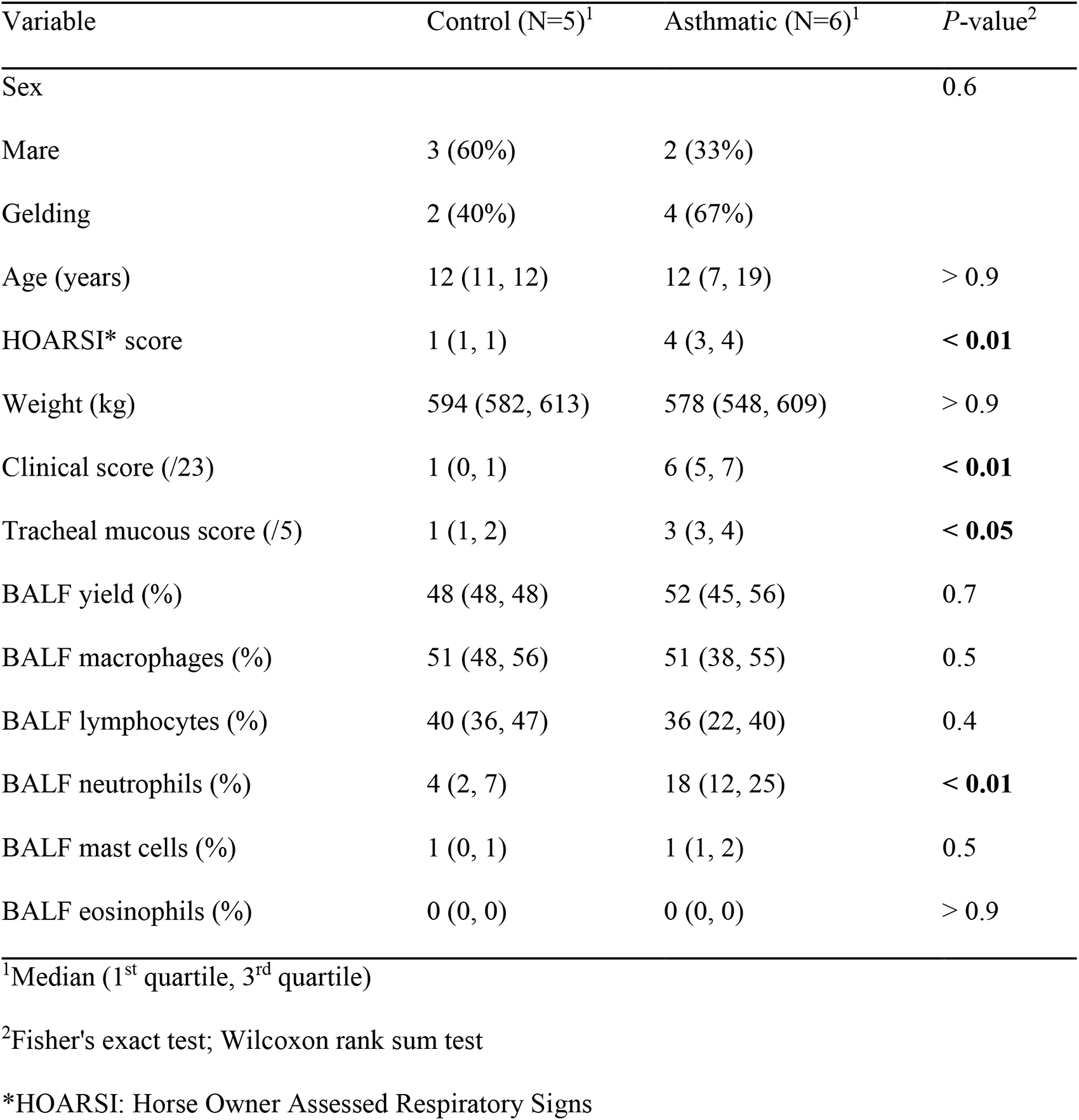
Study population characteristics.

| Variable | Control (N=5)<sup>1</sup> | Asthmatic (N=6)<sup>1</sup> | *P*-value<sup>2</sup> |
| --- | --- | --- | --- |
| Sex |  |  | 0.6 |
| Mare | 3 (60%) | 2 (33%) |  |
| Gelding | 2 (40%) | 4 (67%) |  |
| Age (years) | 12 (11, 12) | 12 (7, 19) | > 0.9 |
| HOARSI* score | 1 (1, 1) | 4 (3, 4) | < **0.01** |
| Weight (kg) | 594 (582, 613) | 578 (548, 609) | > 0.9 |
| Clinical score (/23) | 1 (0, 1) | 6 (5, 7) | < **0.01** |
| Tracheal mucous score (/5) | 1 (1, 2) | 3 (3, 4) | < **0.05** |
| BALF yield (%) | 48 (48, 48) | 52 (45, 56) | 0.7 |
| BALF macrophages (%) | 51 (48, 56) | 51 (38, 55) | 0.5 |
| BALF lymphocytes (%) | 40 (36, 47) | 36 (22, 40) | 0.4 |
| BALF neutrophils (%) | 4 (2, 7) | 18 (12, 25) | < **0.01** |
| BALF mast cells (%) | 1 (0, 1) | 1 (1, 2) | 0.5 |
| BALF eosinophils (%) | 0 (0, 0) | 0 (0, 0) | > 0.9 |

<sup>1</sup>Median (1<sup>st</sup> quartile, 3<sup>rd</sup> quartile)
<sup>2</sup>Fisher's exact test; Wilcoxon rank sum test
\*HOARSI: Horse Owner Assessed Respiratory Signs

Unsupervised clustering of the data identified 19 distinct cell clusters (Fig. 1A). Through automated annotation using the top ten differentially expressed genes (DEGs) derived from major cell types identified in our pilot study (*13*), we successfully predicted the identity of 99.6% of the cells. Cell cluster identities were validated using the expression of known canonical markers and the top DEGs specific to each cell group (Fig. 1B and 1C). Subsequently, the cell clusters were consolidated into six major cells groups: B cells, dendritic cells (DCs), mast cells, monocytes-macrophages (Mo/Ma), neutrophils and T cells. The marker genes used for annotation are compiled in the supplementary tables 2 – 8.

**Figure 1:**
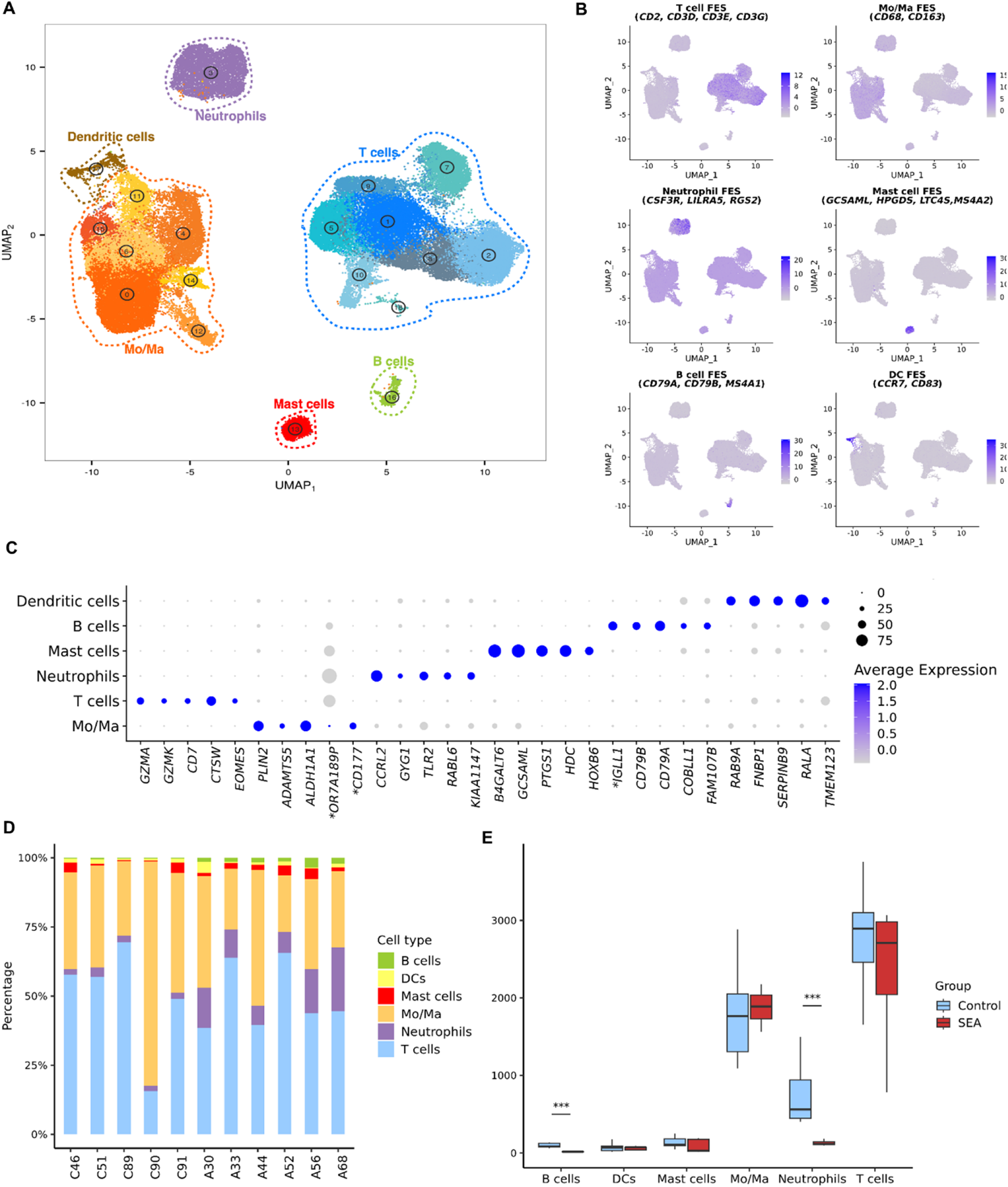
Major cell types identified in the BALF of asthmatic and control horses using scRNA-seq. **(A)** UMAP representation of the 19 clusters identified as six major cell types. *Mo/Ma, monocyte-macrophage.* **(B)** Gene expression patterns of cell type canonical markers. *DC, dendritic cell; FES, feature expression score.* **(C)** Top five differentially expressed genes per major cell type (one non-coding gene removed). *NCBI 103 annotations for *LOC100146200*: *OR7A189P*, *LOC100069985*: *CD177, LOC102147726*: *IGLL1*. **(D)** Distribution of the six major cell types in asthmatic and control horses. **(E)** Number of cells from each major cell type in the asthmatic and control groups. *SEA, severe equine asthma*. ***, *P-*value < 0.001.

### ScRNA-seq reveals the presence of neutrophil subtypes in equine BALF

To explore the diversity of each major cell type, we re-analyzed them independently. Within the neutrophil population, unsupervised clustering revealed three distinct clusters: Neu0, Neu1, and Neu2 (Fig. 2A). The most upregulated gene in Neu0 was *TSC22D3*, a gene involved in neutrophil apoptosis (*14*). Additionally, the mitochondrial gene *ND2*, among the top five DEGs, supported an apoptotic state (Fig. 2C). Upregulated genes involved in neutrophil extracellular trap (NET) formation, including *LOC100054211* (annotated as *H2BC21* in the NCBI database) and *FGL2* (*15*, *16*), further characterized Neu0. Consequently, this cluster was designated as “apoptotic neutrophils”. Neu1 displayed elevated expression of *TREM1*, an established enhancer of pro-inflammatory responses in human and canine neutrophils (*17*). Several pro-inflammatory cytokines genes (*IL1A*, *IL1B*, *CXCL8*, and *CXCL2*) were also overexpressed in this cluster (Fig. 2B). Notably, *G0S2*, *CXCL8*, and *NFKBIA* showed elevated expression levels, which are predictive of septic shock in human peripheral neutrophils (*18*). Thus, this cluster was annotated as “pro-inflammatory neutrophils”. The expression profile of cluster Neu2 was dominated by interferon-stimulated genes (ISGs) such as *IFIT1/2/3/5*, *OAS 2/3* or *ISG15* (*19*), suggesting an anti-viral phenotype (Fig. 2B). A similar gene signature has been observed in circulating neutrophils of healthy humans (*20*) and equine BALF (*21*). Upregulation of the transcription factors *IRF7* and *IRF1*, known to directly activate ISG expression (*19*), further supported the characterization of Neu2 as “ISG^high^ neutrophils”. Our findings confirm the presence of distinct neutrophil subtypes in equine BALF, including pro-inflammatory and ISG^high^ populations previously identified in both healthy and asthmatic horses’ BALF (*21*).

**Figure 2:**
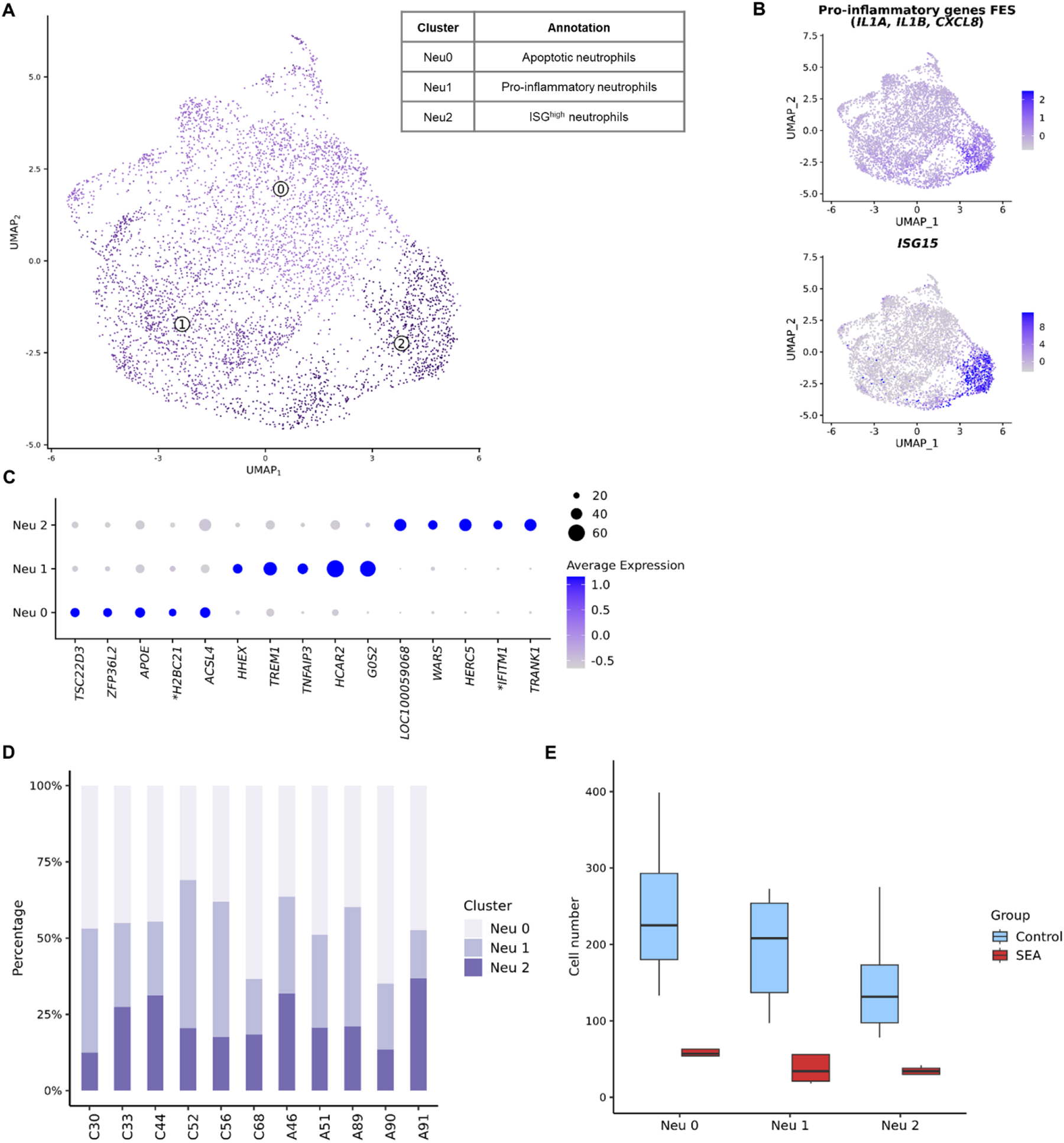
Neutrophils subtypes identified in the BALF of asthmatic and control horses using scRNA-seq. **(A)** UMAP representation of the three clusters identified. **(B)** Gene expression patterns used for annotation. *FES, feature expression score.* **(C)** Top five differentially expressed genes per cluster (one mitochondrial gene removed). NCBI 103 annotation for *LOC100059068*: *IFIT5-like*. *NCBI 103 annotation for *LOC111774805*: *H2BC21*, *LOC100050797*: *IFITM1*. **(D)** Distribution of clusters among asthmatic and control horses. **(E)** Number of cells from each neutrophil cluster in the asthmatic and control groups. *SEA, severe equine asthma*.

### B cell diversity is described for the first time in equine BALF

We identified three distinct B cell clusters through unsupervised clustering: B0, B1, and B2 (Fig. 3A), representing a novel description of B cell diversity in equine BALF. B0 exhibited gene expression patterns consistent with naïve B cells, while B1 and B2 displayed characteristics indicative of plasma (antibody-producing) cells (*13*) (Fig. 3B). B0 was characterized by the upregulation of MHC-II associated genes (*22*) and the downregulation of genes associated with antibody production, such as *TXNDC5*, *HSP90B1*, *TENT5C* and immunoglobulin λ light chain (*IGLL1*)-coding genes. Cluster B1 showed an opposite gene expression pattern compared to B0. Furthermore, genes involved in IgM production (*JCHAIN* and *MZB1*) were upregulated, suggesting that these cells were in the early stages of differentiation into plasma cells. Cluster B2 displayed high levels of galectin 1 (*LGALS1*) mRNAs, a gene upregulated in differentiating plasma cells following immunization and crucial for maintaining antibody secretion (*23*). Therefore, we designated B0 as “naïve B cells”, B1 as “non-switched plasma cells” and B2 as “activated plasma cells”.

**Figure 3:**
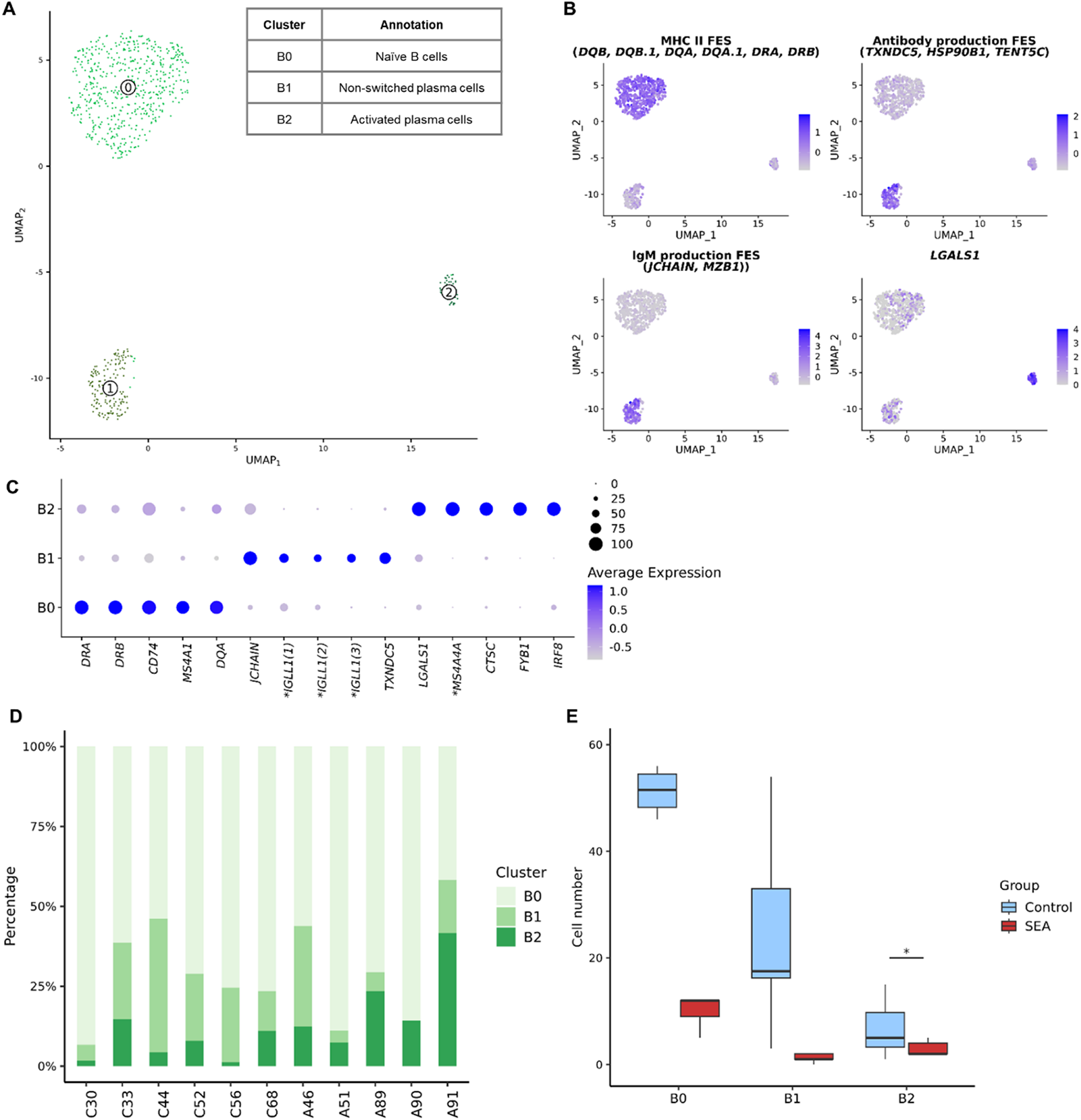
B cell subtypes identified in the BALF of asthmatic and control horses using scRNA-seq. **(A)** UMAP representation of the three clusters identified. **(B)** Gene expression patterns used for annotation. FES, feature expression score. **(C)** Top five differentially expressed genes per cluster. *NCBI 103 annotations for *LOC111774805*, *LOC100060608* and *LOC102147726*: *IGGL1*, for *LOC100061331*: *MS4A4A*. **(D)** Distribution of the clusters among asthmatic and control horses. **(E)** Number of cells from each neutrophil cluster in the asthmatic and control groups. *SEA, severe equine asthma.* \**P-*value < 0.05.

### Equine BALF demonstrates substantial T cell diversity

Unsupervised clustering identified seven T cell clusters (Fig. 4A). The tissue resident marker *ITGAE* was expressed across all T clusters, albeit at lower levels in T6. Clusters T0, T2 and T3 were identified as *CD8*^+^ T cells, while T1 was labeled as “*CD4*^+^ T cells” (T helper cells) based on the differential expression of cell surface markers *CD4*, *CD8a* and *CD8b* (Fig. 4B). Cytotoxicity markers such as *CTSW*, *PRF1*, and *GZMA* were upregulated in T0, T2, and T3. Notably, T0 overexpressed additional cytotoxicity effectors (*NKG7*, *EOMES* and *GZMK*) and the chemokine gene *CCL5*, indicating an “effector CD8^+^ T cells” identity (*24*) (Fig. 4C and 4D). A subset of T0 cells showed upregulation of a *GZMH* isoform, suggesting a higher level of cytotoxic specialization (*25*). Interestingly, some cells in T0 were *CD4*^+^ (Fig. 4B), indicating the presence of a *CD4*^+^ cytotoxic T lymphocytes (*CD4*^+^ CTLs) subset (*26*). Thus, T0 was annotated as “cytotoxic T lymphocytes”.

**Figure 4:**
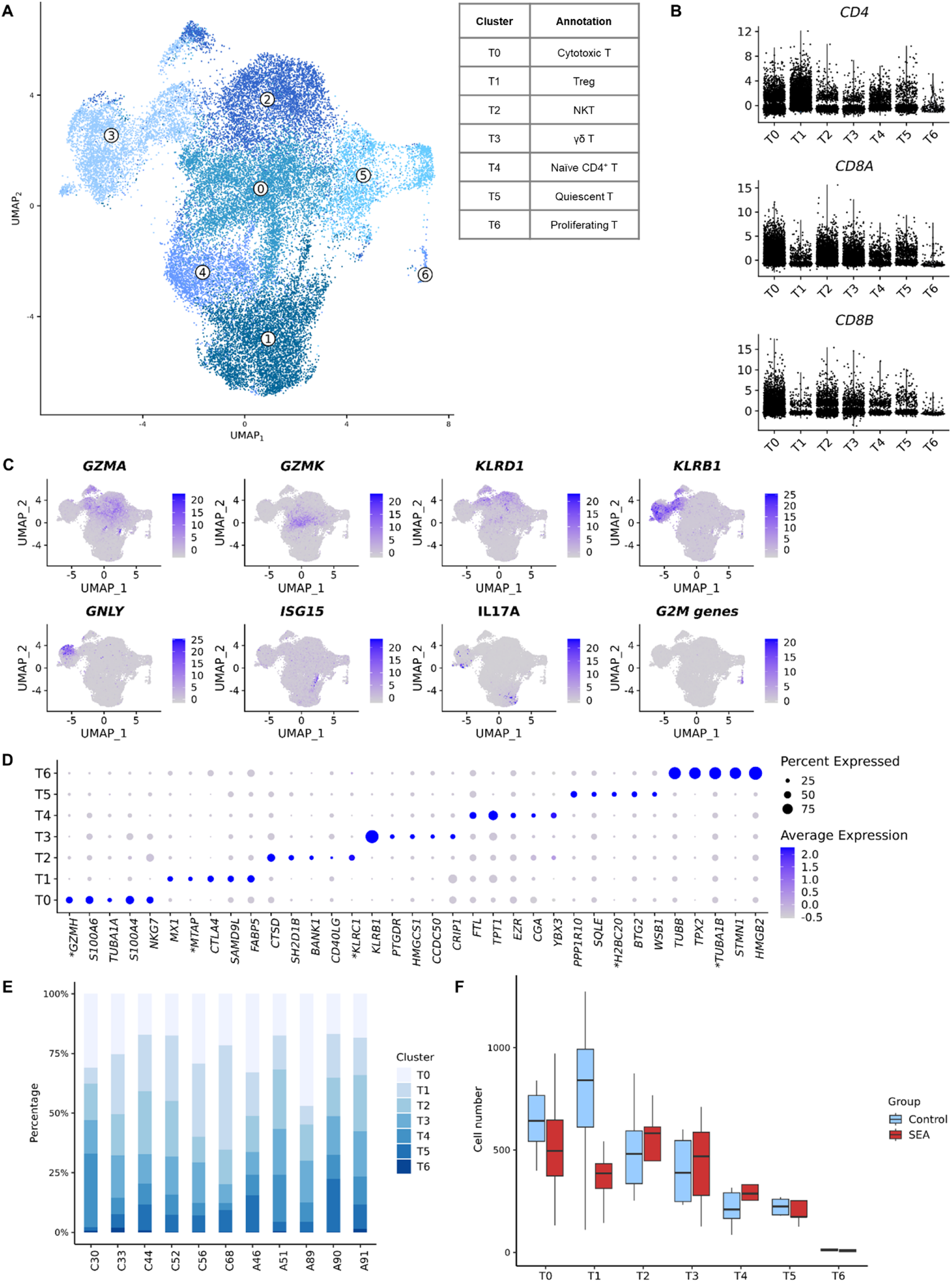
T cell subtypes identified in the BALF of asthmatic and control horses using scRNA-seq. **(A)** UMAP representation of the seven clusters identified. *NKT, Natural Killer T cell.* **(B)** Gene expression patterns used for annotation. **(C)** Top five differentially expressed genes per cluster (snRNA, non-coding genes and ribosomal protein genes removed). *NCBI 103 annotations for *LOC100051986*: *GZMH*, *LOC100065392*: *MTAP*, *LOC100062823*: *KLRC1*, *LOC100053968*: *H2BC20*, *LOC100059091*: *TUBA1B*. **(D)** Expression levels of the cell surface markers *CD4*, *CD8a* and *CD8b*. **(E)** Distribution of the clusters among asthmatic and control horses. **(F)** Number of cells from each T cell cluster in the asthmatic and control groups. *SEA, severe equine asthma*.

The expression profile of T2 resembled that of previously identified natural killer T (NKT) cells in equine BALF (*13*), displaying both natural killer (NK)-specific (*SH2D1B*, *KLRC1* and *KLRD1*) (Fig. 4D) and T cell-specific (*TRAT1*) features. *TYROBP*, a marker for NKT cells in human peripheral blood, was also upregulated (*27*). T3 was identified as “γδ T cells” based on the upregulation of *KLRB1* (Fig. 4D) and the gene coding for a SCART1-like protein (*13*). This cluster comprised a subset of cytolytic cells, as shown by the expression of *GNLY,* which codes for granulysin (*28*) (Fig. 4C).

T1, labeled as “regulatory T cells” (Treg), exhibited a *CD4*^+^*FOXP3*^+^ phenotype. Top DEGs included Treg-specific genes such as *CTLA4*, *TRIB1*, *IL32*, *FGL2* or *FOXO1*. Within T1, a subset displayed an ISG^high^ signature previously associated with Tregs (*29*), while another subset showed overexpression of the pro-inflammatory cytokine *IL17A*, which can be induced in Tregs under inflammatory conditions (*30*) (Fig. 4C). Cluster T4 exhibited unspecific DEGs, making annotation challenging. We could not ascertain whether the cluster was *CD4*^+^*CD8*^-^ or double negative *CD4*^-^ *CD8*^-^ on the violin plots (Fig. 4B). Based on the absence of upregulation of canonical markers for double negative T cells (*31*), we presumed these cells to be *CD4*^+^*CD8*^-^. *LEF1* and *CCR7*, typically expressed by naïve T cells, were upregulated, and ribosomal protein genes were overexpressed, indicating an ongoing differentiation process (*32*) (Suppl. fig. 1). Cluster T4 was thus annotated as “naïve CD4^+^ T cells”. Cells in T5 demonstrated upregulation of genes linked to the S phase (e.g., *BTG2*, *PLK2* and *MCL1*) and downregulation of ribosomal protein genes, suggesting a quiescent state. Conversely, T6 cells exhibited high levels of mitosis markers, indicating proliferating cells (Fig. 4C). T5 and T6 were probably composed of both *CD4*^+^ and *CD8*^+^ phenotypes, as there was no clear increase in the expression of *CD4* or *CD8* genes in these clusters.

### The monocyte-macrophage population is constituted from monocytes, several alveolar macrophages subtypes, and putative monocyte-lymphocyte complexes

Monocyte-macrophage diversity was explored through unsupervised clustering, revealing six clusters (Fig. 5A). Clusters Mo/Ma0, Mo/Ma1, Mo/Ma2 and Mo/Ma5 were identified as alveolar macrophages (AMs) based on specific marker expression (Fig. 5B). Clusters Mo/Ma3 and Mo/Ma4 were annotated as monocytes.

**Figure 5:**
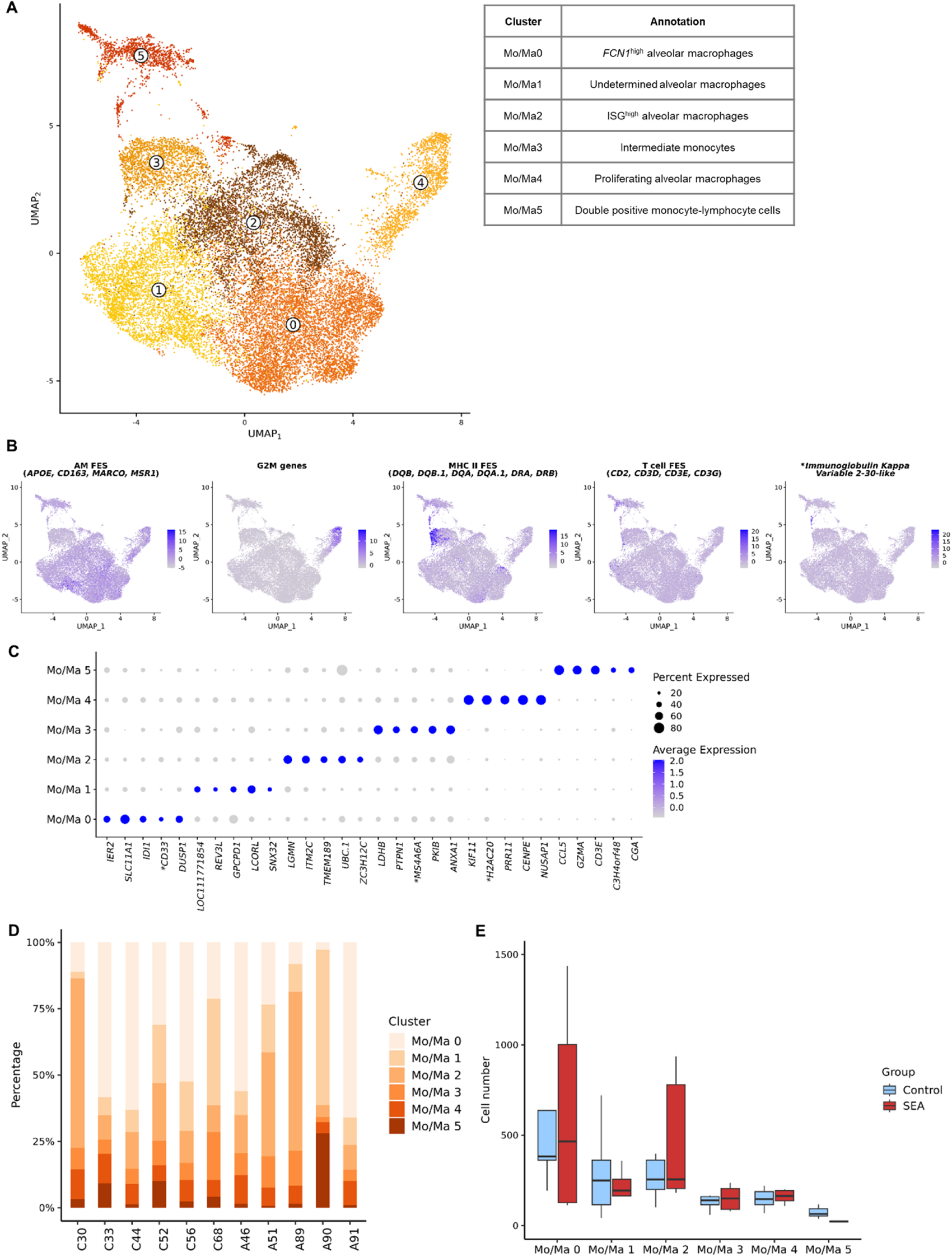
Monocytes-macrophages (Mo/Ma) subtypes identified in the BALF of asthmatic and control horses using scRNA-seq. **(A)** UMAP representation of the six clusters identified. **(B)** Gene expression patterns used for annotation. *FES, feature expression score.* *NCBI 103 annotations for *LOC100147522*: *GZMH.* **(C)** Top five differentially expressed genes per cluster (non-coding, mitochondrial and ribosomal protein genes removed). NCBI 103 annotation for *LOC111771854*: *oleosin-B6-like*. *NCBI 103 annotations for *LOC100066849*: *CD33*, *LOC100061154*: *MS4A6A, LOC100058587*: *H2AC20*. **(D)** Distribution of the clusters among asthmatic and control horses. **(E)** Number of cells from each Mo/Ma cluster in the asthmatic and control groups. *SEA, severe equine asthma*.

Mo/Ma0 displayed an anti-inflammatory phenotype, with upregulation of *DUSP1*, *LILRB4*, *IRS2* genes and downregulation of *NFKB1* and *NFKB2* genes. This cluster resembled the “*FCN1*^high^ AM” previously identified in equine BALF (*13*), with overexpression of *FCN1* (*LOC100069029*) and *ORM2* (*LOC100050034* and *LOC100050100* isoforms). Mo/Ma1 had a lower feature count, a higher percentage of mitochondrial reads (Suppl. fig. 2), and no specific gene expression pattern related to a particular cell type or function. It could potentially represent either fragile or quiescent cells. Genes associated with cell adherence (*CDH23*, *RASAL2*) were upregulated, supporting a population of quiescent resident AMs. This cluster was conservatively annotated as “undetermined AM”. Mo/Ma2 showed upregulation of *LGMN* and *UBC*, associated with a pro-inflammatory (M1) phenotype (*33*, *34*), as well as a predominant expression of ISGs (e.g., *OASL*, *IFI6*, *IFI44* or *IRF7*). We labeled the cluster “ISG^high^ AM”. Mo/Ma4 displayed a gene expression profile indicative of “proliferating AMs” (Fig. 5B).

Mo/Ma3 exhibited a similar expression profile to intermediate monocytes in our pilot study (*13*), with upregulation of genes *SPP1*, *CCL15*, *CD44* and *MMP9*, as well as high levels of MHCII-associated genes (Fig. 5B). This cluster was thus labeled as “intermediate monocytes”. The overexpression of genes associated with antigen processing and presentation (*IFI30*, *TGFB1*, *CTSB* and *CD74*) suggested ongoing maturation into macrophages. Their relatively high expression of ribosomal protein genes supported this hypothesis, as these genes are typically upregulated during early differentiation (*32*) (Suppl. fig. 3).

Mo/Ma5 showed a distinct T cell signature with the expression of *CD2*, *CD3D*, *CD3E*, *CD3G* and *CD7*, among others (Fig. 5B). A subset of Mo/Ma5 exhibited a B cell signature, with overexpression of *LOC100630729*, encoding an Igκ-like protein (Fig. 5B). The gene expression pattern of Mo/Ma5 resembled the double positive monocyte-lymphocyte cells we previously identified in equine BALF (*13*). We hypothesized that this cluster represents immune cell complexes similar to those found in human peripheral blood (*35*, *36*). Mo/Ma5 also demonstrated overexpression of MHCII-associated genes, critical for immunological synapse formation (Fig. 5B).

### Distinct clusters of dendritic cells are present in equine BALF

Unsupervised clustering identified four distinct DC clusters: DC0, DC1, DC2 and DC3 (Fig. 6A). DC0, characterized by the upregulation of *CD1C*, *CLEC10A, FCER1A* (*37*) and MHCII-associated genes (*38*), was annotated as conventional DC2 (cDC2) (Fig. 6C). DC1 showed an activation profile with upregulated genes from a previously described DC activation panel, including *CCR7*, *LAMP3*, *IDO1*, *FSCN1* and *CD83* (*37*) (Fig. 6C). *SERPINB9*, a marker for cross-presentation capable DCs (*39*), was upregulated in this cluster, along with downregulation of MHCII-associated genes (*40*), indicating a mature DC phenotype (Fig. 6C). DC1 was annotated as “*CCR7*^+^ DCs”, which have been also referred to as “activated DCs” or “mature DCs enriched in immunoregulatory molecules” (*37*). In humans, several DC types can display this activation profile. Our equine *CCR7*^+^ DCs population may thus similarly be composed from different lineages.

**Figure 6:**
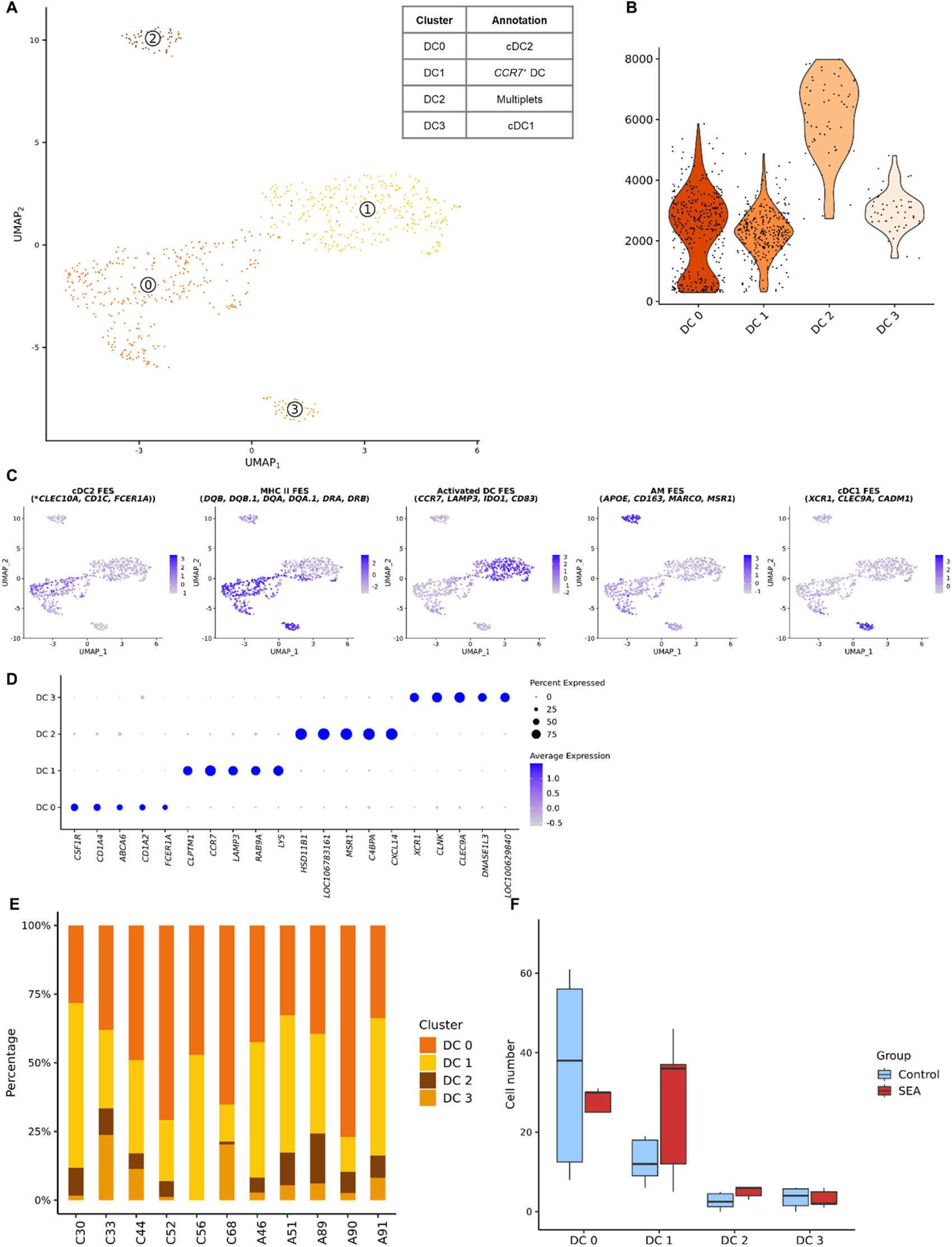
Dendritic cell (DC) subtypes identified in the BALF of asthmatic and control horses using scRNA-seq. **(A)** UMAP representation of the four clusters identified. *cDC, conventional dendritic cell.* **(B)** Gene expression patterns used for annotation. *FES, feature expression score*. *NCBI 103 annotations for *LOC100072936* and *LOC100072933*: *CLEC10*. **(C)** Top five differentially expressed genes per cluster (non-coding genes removed). NCBI 103 annotations for *LOC100629840*: *bone marrow proteoglycan-like*, *LOC106783161*: *apolipoprotein R*. **(D)** RNA feature count for each DC cluster. **(E)** Distribution of the clusters among asthmatic and control horses. **(F)** Number of cells from each DC cluster in the asthmatic and control groups. *SEA, severe equine asthma*.

Cluster DC2 showed upregulated markers for AMs (*APOE*, *CD163*, *MARCO* and *MSR1*), cDCs (*CXCL13*, *SIRPA*) and plasmacytoid DCs (*IRF7*) (Fig. 6C). The high RNA feature count in this cluster further supported the presence of doublets and/or multiplets, and it was thus annotated as “multiplets” (Fig. 6B) and was not used for further analysis. DC3 was annotated as “cDC1” based on the expression of *XCR1*, *CLEC9A* and *CADM1* (*37*) (Fig. 6C).

### The mast cells form a homogenous cluster in our study population

The mast cells formed a highly homogenous cell cluster, without any convincing subclusters identified in independent reanalysis. This finding contrasts with a previous scRNA-seq study on the BALF from asthmatic and control horses, in which four distinct mast cell clusters were identified (*21*). This study included horses affected by a different type of equine asthma, a milder form characterized by a high proportion of mast cells in the BALF. The larger number of mast cells in their study likely facilitated the identification of subtle differences between subtypes.

### Sample processing and counting techniques do not significantly influence cellular composition

To evaluate the potential impact of sample processing and cryopreservation on cellular composition, we compared the distribution of the five cytologically distinguishable major leukocytes (lymphocytes, macrophages, neutrophils, mast cells, and eosinophils). While some variations were observed among individual horses, as shown in supplementary figure 4, repeated-measure analysis of variance (ANOVA) did not reveal any significant differences, indicating that sample processing did not have a substantial effect on the cellular composition of the samples.

Similarly, we compared the proportions of each cell type obtained through manual cell counting on cell suspension cytology with those obtained through scRNA-seq. Once again, the repeated-measure ANOVA did not detect any significant differences.

### The BALF of asthmatic horses is enriched in B cells but specifically depleted in activated plasma cells

As expected, asthmatic horses showed a significantly higher proportion of neutrophils compared to the control horses (Table 2, fig. 1D and 1E). A novel finding was the B cell enrichment in the BALF of asthmatic horses (Fig. 1D and 1E). Upon analyzing the distribution of B cell subtypes between groups, we found that asthmatic horses exhibited approximately three times fewer activated plasma cells (B2 cluster) than control horses (Table 2, fig. 3D and 3E). This suggests that expansion of the naïve B cells and non-switched plasma cells primarily contributed to the increased B cell proportion in asthmatic horses. No significant differences were observed between asthmatic and control groups for other major cell types or subtypes (Table 2 and fig.1E, 2E, 4F, 5E, 6F).

**Table 2:**
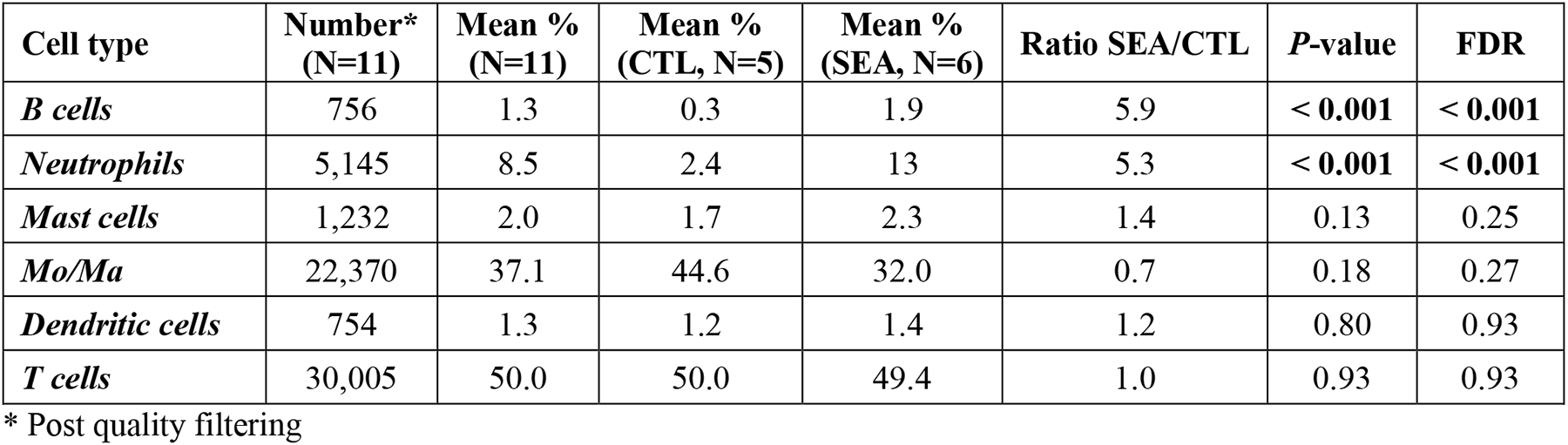
Proportions of the major cell types determined with scRNA-seq and compared between asthmatic (SEA) and control (CTL) groups.

| Cell type | Number*<br>(N=11) | Mean %<br>(N=11) | Mean %<br>(CTL, N=5) | Mean %<br>(SEA, N=6) | Ratio SEA/CTL | P-value | FDR |
| --- | --- | --- | --- | --- | --- | --- | --- |
| *B cells* | 756 | 1.3 | 0.3 | 1.9 | 5.9 | < 0.001 | < 0.001 |
| *Neutrophils* | 5,145 | 8.5 | 2.4 | 13 | 5.3 | < 0.001 | < 0.001 |
| *Mast cells* | 1,232 | 2.0 | 1.7 | 2.3 | 1.4 | 0.13 | 0.25 |
| *Mo/Ma* | 22,370 | 37.1 | 44.6 | 32.0 | 0.7 | 0.18 | 0.27 |
| *Dendritic cells* | 754 | 1.3 | 1.2 | 1.4 | 1.2 | 0.80 | 0.93 |
| *T cells* | 30,005 | 50.0 | 50.0 | 49.4 | 1.0 | 0.93 | 0.93 |

\* Post quality filtering

**Table 3:**
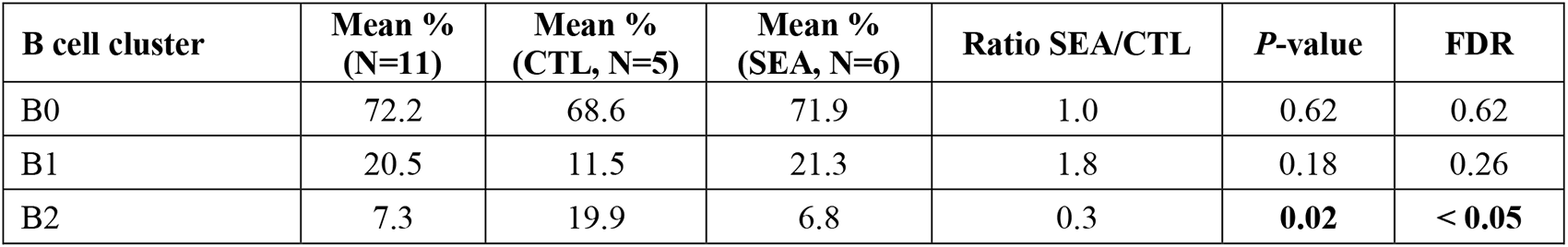
Proportions of the B cell subtypes identified with scRNA-seq and compared between asthmatic (SEA) and control (CTL) groups.

| B cell cluster | Mean %<br>(N=11) | Mean %<br>(CTL, N=5) | Mean %<br>(SEA, N=6) | Ratio SEA/CTL | P-value | FDR |
| --- | --- | --- | --- | --- | --- | --- |
| B0 | 72.2 | 68.6 | 71.9 | 1.0 | 0.62 | 0.62 |
| B1 | 20.5 | 11.5 | 21.3 | 1.8 | 0.18 | 0.26 |
| B2 | 7.3 | 19.9 | 6.8 | 0.3 | **0.02** | < **0.05** |

### Gene expression profile of neutrophils indicates altered NETosis and migratory function in SEA

Using a mixed model approach, we compared gene expression between asthmatic and control horses within each cell type and subtype. Supplementary tables 9 – 13 provide the results of the differential gene expression (DGE) analysis, where positive log fold change indicates upregulation in the SEA group.

In asthmatic horses, neutrophils showed significant changes in gene expression. We observed 13 upregulated genes and 206 downregulated genes in this cell population. The asthmatic neutrophils exhibited an “asthma signature” characterized by upregulation of *CHI3L1* and *MAPK13*, known markers of neutrophilic asthma in humans (*41–43*), and downregulation of *SLC7A11*, an indicator of ferroptosis reduced in neutrophilic mice asthma (*44*). Apoptotic neutrophils (Neu0) upregulated *S100A9* and *RETN*, both involved in NETosis function. NETosis is a form of neutrophil death in which decondensed chromatin and granular contents (so-called NETs) are released into the extracellular space. *S100A9* is a key mediator of neutrophilic asthma and plays a role in neutrophil activation and NETs formation (*45*, *46*). *RETN*, which codes for resistin, is associated with increased susceptibility of neutrophils to LPS activation, and to enhanced NETosis (*47*). Pro-inflammatory neutrophils (Neu1) of asthmatic horses only upregulated one gene, *PTGS2*, and downregulated six genes, including *KLF2*. Reduced *KLF2* levels can promote neutrophil migration (*48*) and exacerbate NET-induced transfusion-related acute lung injury (*49*). In the ISG^high^ neutrophils (Neu2), we observed upregulation of *GIBp6*, an ISG, and *ADGRE5*, also known as *CD97*. *ADGRE5* activation may promote migration of ISG^high^ neutrophils to the lungs (*50*).

Interestingly, we also identified gene expression features that could have a protective effect on the lower airways. The antileukoproteinase gene *SLPI* was upregulated in asthmatic horses, which can provide anti-inflammatory actions by inhibiting the NFκB pathway and preventing excess NET formation (*51*). *NFKB1* was indeed downregulated in asthmatic neutrophils. Downregulation of *IL17RC* suggested a reduced capacity to respond to the Th17 cytokine IL17. The predominant contributor of protective features was the apoptotic neutrophil subtype, with upregulation of *SLPI* and downregulation of *CCL20* and *NR4A3*. The Th17-associated cytokine CCL20 is a potent chemotactic factor for lymphocytes and DCs. The downregulation of *CCL20* could thus have an overall anti-inflammatory effect, with reduced chemotaxis of immune cells and reduced Th17-signalling. *NR4A3* positively regulates neutrophil survival (*52*). Hence, its downregulation may mitigate neutrophil persistence in the lungs in SEA.

In summary, neutrophils from asthmatic horses exhibited DGE patterns indicative of asthma, including known markers of human asthma. Moreover, these neutrophils displayed an expression profile consistent with increased migratory capacity and the potential for NET formation. The co-expression of genes with protective roles suggests a dual pro- and anti-inflammatory role of neutrophils in SEA.

### Upregulation of POU2AF1 in asthmatic B cells reveals potential links to pulmonary fibrosis

The DGE analysis conducted on B cells identified five DEGs in asthmatic horses. The only upregulated gene was *POU2AF1*, a transcriptional coactivator essential for B cell function. Elevated expression of *POU2AF1* in B lymphocytes has been associated with interstitial pulmonary fibrosis (*53*) and chronic obstructive pulmonary disease (COPD) (*54*) in humans. Notably, its expression has shown a negative correlation with lung function (*54*). The DGE analysis between B cell subtypes did not yield any significant DEGs.

### Gene expression patterns of asthmatic T cells support a Th17-oriented immune response

The DGE analysis in T cells revealed 77 upregulated genes and four downregulated genes in asthmatic horses. Notably, two known markers of human asthma, *IL26* (*55*) and *OLFM4* (*56*), were upregulated. As in neutrophils, the acute asthma marker *RETN* (*57*) was upregulated in cytotoxic T cells (T0). Moreover, T cells from asthmatic horses exhibited a robust Th17 signature, characterized by the upregulation of *IL17A*, *IL17F*, *IL21* and *CCL20*.

Interestingly, naïve *CD4*^+^ T cells (T4) showed a simultaneous upregulation of Th17-associated genes (*IL17A*, *IL1B*, *CCL20* and *NFKBID*) and *FOXP3*. This observation supported the hypothesis that naïve *CD4*^+^ T cells in asthmatic horses adopt a Th17 pathway during differentiation, as *FOXP3* expression is transiently present during Th17 cell development (*58*).

Furthermore, the Treg cells (T1) displayed a Th17-oriented profile, with upregulation of *IL21* and *IL17A* and downregulation of *EOMES*, a known suppressor of Th17 differentiation in human Treg cells (*59*) (Fig. 7).

**Figure 7:**
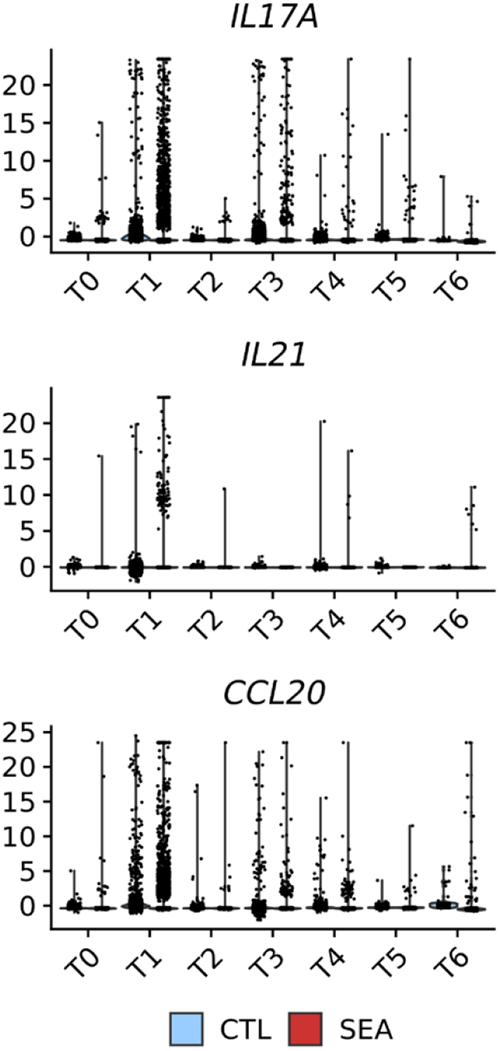
Upregulation of Th17-associated genes in the T cells of severely asthmatic horses (SEA, right) compared to control horses (CTL, left).

The γδ T (T3) cells conjointly upregulated *IL17A* and *IL1R*, consistent with a γδ17 phenotype (*60*). In mice, γδ T cells possess an intrinsic capacity for IL17 production, which is directly induced by IL23 and IL1 (*61*). Notably, γδ17 T cells are implicated in various human inflammatory diseases (*61*), and increased *IL1R* expression has been associated with neutrophilic asthma and reduced pulmonary function in humans (*60*).

### Genes associated with T cell function are dysregulated in SEA

Several genes involved in T cell function were differentially expressed in asthmatic horses. Specifically, the marker of T cell exhaustion, *TOX2* (*62*), was upregulated, along with *S1PR5*, whose expression is induced by antigen exposure (*63*). Cytotoxic T cells (T0) downregulated *GZMB*, a gene associated with lymphocytic inflammation in the lungs (*64*). The downregulation of *IL18R1* and *XCL1* supported Treg cell dysfunction. Indeed, downregulation of the IL18 receptor is associated with unresponsiveness of exhausted *CD8*^+^ T cells (*65*). Furthermore, dysfunctional Treg cells in individuals with allergic asthma have been shown to downregulate *XCL1* (*66*). Among the T cell subtypes, NKT cells (T2) upregulated *NPY*, a gene associated with reduced NK function (*67*).

Surprisingly, the T cells and several T cell subtypes upregulated a few B cell-specific immunoglobulin genes, including *LOC102147726* (coding for the immunoglobulin λ1 light chain *IGLL1*), *LOC100060608* (coding for the immunoglobulin λ constant 7 *IGLC7*), and *JCHAIN*. To investigate this further, we examined the absolute gene expression level in T cells and found that the log counts of immunoglobulin genes were three to seven times lower than the mean gene expression level in the T cell cluster. Assuming a spurious signal, we concluded that the upregulation of immunoglobulins was unlikely to have biological relevance in our dataset.

### Monocytes and alveolar macrophages from asthmatic horses display a Th17 signature

The DGE analysis of Mo/Ma major cell type identified 35 upregulated genes and four downregulated genes in the asthmatic group. Among the upregulated genes were *OLFM4*, associated with severe lung disease in humans (*68*, *69*), and the marker of neutrophilic asthma *CHI3L1* (*42*). *S100A8*, shown for its increased expression in individuals with steroid-resistant neutrophilic asthma, was upregulated (*70*). *TLR1*, recently identified as a potential therapeutic target for asthma in humans (*71*), was also differentially expressed.

In the *FCN1*^high^ AMs (Mo/Ma0), positively DEGs included *PGLYRP1*, *PTX3* and *CCL20*. In mice, *PGLYRP1* promotes pro-asthmatic Th2 and Th17 responses (*72*), while *PTX3* is a marker of non-eosinophilic asthma in humans (*73*). In horses, BALF *PTX3* expression increases in acute asthmatic crisis, particularly in dust-activated foamy macrophages (*74*). The simultaneous upregulation of the Th17-associated cytokine *CCL20* and downregulation of the Th1-cytokine *CCL11* supported a Th17 polarization of *FCN1*^high^ AMs in asthmatic horses. Moreover, in the ISG^high^ AMs (Mo/Ma2), genes encoding *CCL20* and its receptor *CCR6* were upregulated, further advocating for a Th17 phenotype.

An important finding was the significant upregulation of *CXCL13* in Mo/Ma, intermediate monocytes (Mo/Ma3), and putative monocyte-lymphocyte complexes (Mo/Ma5). This B cell chemoattractant has been previously shown to be upregulated in peripheral blood cells from horses with SEA, particularly after stimulation with hay dust extract (*75*). *CXCL13* cytokine levels are also elevated in the serum and the BALF of asthmatic humans (*76*, *77*). In an OVA murine model of asthma, an anti-CXCL13 antibody reduced cell recruitment, bronchial-associated lymphoid tissue formation, and airway inflammation, highlighting *CXCL13* as a promising therapeutic target (*76*). Macrophages and Th17-derived cells are the primary sources of *CXCL13* production. In this single-cell dataset, *CXCL13* was found to be upregulated in intermediate monocytes and putative monocyte-lymphocyte complexes from asthmatic horses, but not in alveolar macrophages or T cells. These findings suggest that the observed increase in *CXCL13* in horses and humans likely results from activated monocytes’ upregulation of *CXCL13*, rather than expansion of macrophage or Th17 populations.

Furthermore, intermediate monocytes in asthmatic horses demonstrated upregulation of *S100A9*, *S100A12*, *CCL17* and *S1PR5*. *S100A9* and *S100A12* serve as biomarkers for neutrophilic asthma (*78*, *79*), while *CCL17* is associated with asthma and may contribute to airway remodeling through fibroblast activation via the CCR4-CCL17 axis (*80*, *81*). Intriguingly, *S1PR5* regulates monocyte trafficking (*82*), suggesting intermediate monocytes from asthmatic horses may possess a higher migratory capacity.

In contrast to the predominant Th17 signature observed in the Mo/Ma clusters, we noted upregulation of the Th2-associated genes *PRB1* and *NMUR1* in the Mo/Ma major cell population (*83*, *84*).

### Th17 activation may result from a crosstalk between monocytes and lymphocytes

The presence of multiple cell types within the Mo/Ma5 cluster was supported by the high number of DEGs identified. This cluster exhibited simultaneous upregulation of *CXCL13* and *IL17A*, both associated with the Th17 pathway. Interestingly, while several T cell clusters in the dataset upregulated *IL17A*, none of the Mo/Ma clusters, except Mo/Ma5, showed this upregulation. Conversely, *CXCL13* upregulation was exclusive to Mo/Ma5 and not observed in any T cell clusters. This led us to conclude that the co-upregulation of *IL17A* and *CXCL13* originated from the dual nature of Mo/Ma5 as monocyte-lymphocyte complexes. Downregulation of the Th1-associated gene *CD27* and granzyme B-like proteins further suggested a Th17 polarization within the cells composing the complexes (*85*). Additionally, upregulation of inflammasome-related genes (*SIGLEC14*, *KCNK13* and *PELI2*) (*86*) was observed.

Similar to T cells, upregulation of B-cell specific genes (*IGLL1* and *IGLC7*) was observed in multiple Mo/Ma clusters. However, their average log counts were considerably lower compared to the mean gene expression level in Mo/Ma, rendering their upregulation irrelevant. On the other hand, *JCHAIN* was upregulated specifically in Mo/Ma1 (undetermined AMs), and its absolute expression level closely aligned with the mean gene expression within the cluster. Possible explanations include technical doublets, *bona fide* AM-B cell complexes, phagocytized B cells, or the expression of immunoglobulin genes by activated AMs themselves, as recently demonstrated in human and mice (*87*).

### Gene expression patterns of asthmatic DCs suggest enhanced migratory capacity and non- Th2 response

The DGE analysis of DCs identified 14 DEGs, of which nine were upregulated in asthmatic horses. None of the upregulated genes had been previously associated with asthma or dendritic cell function. On the other hand, downregulated genes included *MARCO* and *RSAD2*. In a murine OVA-asthma model, *MARCO*-deficient mice showed increased eosinophilic airway inflammation and airway hyperresponsiveness, accompanied by enhanced migration of lung DCs to draining lymph nodes (*88*). Consequently, reduced *MARCO* expression in equine lung DCs may enhance their migration to lymph nodes, leading to an amplified immune response against aeroallergens.

Further analysis of DC subtypes yielded significant results for DC0 (annotated as cDC2s) only, with one upregulated (*GLRX2*) and five downregulated genes. Administration of GLRX2 has been shown to reduce airway inflammation in an OVA-asthma model (*89*), indicating its potential protective function. The downregulation of the chemokine gene *CCL8* was of particular interest. CCL8 is responsible for the recruitment of basophils, eosinophils and mast cells in allergic processes and contributes to airway allergic inflammation by promoting a Th2 immune response (*90*). Hence, *CCL8* downregulation in cDC2s suggests a shift towards a non-Th2 response in SEA.

### Downregulation of *YBX3* in mast cells of asthmatic horses points to potential airway remodeling

In mast cells, eight DEGs were identified, with two (*LYZ* and *PTPRB*) upregulated in asthmatic horses. Six genes were downregulated, including *YBX3*. Reduced circulating *YBX3* mRNA is a sensitive predictor of idiopathic pulmonary fibrosis in humans (*91*). Therefore, the downregulation of *YBX3* in mast cells of asthmatic horses may contribute to the observed airway remodeling seen in SEA.

## DISCUSSION

Severe equine asthma is characterized by neutrophilic inflammation in the lower airways, resembling a subset of non-Th2 asthma observed in humans. In this study, we utilized scRNA-seq to investigate the immune mechanisms underlying SEA. Among the six major cell types identified, B cells and neutrophils were more abundant in asthmatic horses. Notably, the fraction of activated (switched) plasma cells was decreased, indicating a non-Th2 response. Both T cells and Mo/Ma displayed a strong Th17 signature, including upregulation of *CXCL13* by intermediate monocytes. Furthermore, a subset of cells exhibited an expression profile indicative of monocyte-lymphocyte complexes, potentially contributing to Th17 activation. Neutrophils appeared as bystanders of the lung inflammatory response, with an increase in NETosis function and reduced capacity to respond to Th17 signals. These findings support a primary Th17-mediated immune response in neutrophilic SEA of horses, potentially initiated through crosstalk between monocytes and T cells via direct contact.

Similar Th17-associated responses have been observed in non-Th2 asthma in humans, including organic dust-induced asthma and a subset of non-Th2 asthma patients (*7*, *92*). In horses, previous studies have also implicated the Th17 response in SEA. Increased levels of *IL17* mRNA have been observed in the BALF of horses with SEA following antigen challenge (*9*). Dysregulation of miRNA in the serum of asthmatic horses supported the existence of a mixed Th2/Th17 response (*10*). Furthermore, a comprehensive miRNA-mRNA study in equine lung tissues have suggested a predominant Th17 pathway, along with some indications of a parallel Th2-type response (*11*). Transcriptomics, proteomics, and tissue staining analyses of mediastinal lymph nodes in horses have further supported a predominant Th17 response in severe equine asthma (*12*).

While studies on asthma in mice and humans have mainly focused on T cells as the main contributors to the Th17 response (*92*), our study demonstrated the involvement of both T cells and Mo/Ma populations in driving the Th17 response in horses with asthma. Importantly, this resulted from alterations in gene expression patterns rather than expansion of these cell populations. The upregulation of key Th17 cytokines such as *IL17A*, *IL21*, and *CCL20* was observed in T cell clusters, suggesting their engagement in a Th17 differentiation pathway. Alveolar macrophages and intermediate monocytes also exhibited a strong Th17 signature, with upregulation of *CXCL13*, *PGLYRP1*, *CCL20* or *CCR6*. *CXCL13,* a B cell chemoattractant predominantly produced by Mo/Ma and Th17-derived cells (*93*), has been shown to be upregulated in hay dust extract-stimulated PBMCs of asthmatic horses (*75*). Because the latter study was performed on a cell mixture, the cellular origin of *CXCL13* could not be ascertained. Our findings confirm that *CXCL13* in asthmatic horses mainly originates from activated monocytes. The release of IL17 by T cells probably induces *CXCL13* upregulation in equine lung monocytes, as demonstrated in asthmatic individuals (*94*). Activated monocytes could in turn induce Th17 differentiation of T cells (*95–97*). Collectively, our results support a crosstalk between *IL17A*-producing T cells and *CXCL13*-producing monocytes in the context of Th17-mediated immune response in SEA.

Of particular interest was the double positive cluster Mo/Ma5, which displayed a similar transcriptomic profile as previously observed in equine BALF scRNA-seq studies (*13*, *21*). The presence of monocyte-T cell interactions has been reported in human blood, with the frequency and phenotype of these cell-cell complexes varying depending on the immune response polarization (*35*). Considering that the crosstalk between monocytes and T cells plays a key role in the development of various human inflammatory diseases (*95–97*), the potential presence of *bona fide* monocyte-lymphocyte complexes in the lower airway compartment is particularly intriguing. The reciprocal activation of monocytes and lymphocytes may occur through direct cellular contact rather than solely through endocrine or paracrine mechanisms.

In contrast to previous reports (reviewed in (*2*, *3*, *8*)), we did not detect a Th2 or Th1 signature in the cells from asthmatic horses, except for the upregulation of the Th2-associated genes *PRB1* and *NMUR1* in Mo/Ma. Notably, we did not observe upregulation of characteristic Th2 and Th1 cytokines such as *IL4*, *IL13* or *IFNγ*. Consistent with our findings, Th2 and Th17-associated gene expression appears to be regulated in opposite direction in the airways of human patients (*98*). In horses affected by SEA, downregulation of *IL4* has been shown to correlate with increased IL17 staining intensity in the mediastinal lymph nodes (*12*). Moreover, Th1-and Th2-associated genes are downregulated in antigen-challenged PBMCs from asthmatic horses (*75*). An additional argument against a Th2 response is the reduced fraction of activated plasma cells in asthmatic horses within our study population. Upon antigen stimulation, non-switched IgM-producing plasma cells become activated and produce immunoglobulins of other classes, which is a prerequisite for the Th2 response. While switched plasma cells were less frequent in asthmatic horses, the proportion of total B and plasma cells was significantly higher, likely due to *CXCL13* signaling (*94*). Consequently, asthmatic horses have a larger pool of B cells, which can potentially differentiate into plasma cells and be activated. This could explain the increased susceptibility of asthmatic horses to certain Th2-associated diseases, such as insect bite hypersensitivity and urticaria (*99*). Overall, our findings indicate that SEA is primarily driven by a Th17-mediated immune response characterized by an *IL17*-induced *CXCL13*-mediated recruitment of B cells into the lower airways. The resulting increase in B cell abundance may predispose asthmatic horses to secondary Th2-type responses.

The transcriptomic profile of T cells suggested alterations in T cell function, including T cell exhaustion, unresponsiveness of Treg cells, and reduced cytotoxicity in NKT cells. It remains unclear whether these dysregulations are associated with the Th17 polarization of the T cell populations, or if they represent independent mechanisms. Nevertheless, these alterations in T cell function may potentiate the abnormal immune response observed in SEA.

Neutrophils are short-lived cells that undergo apoptosis rapidly after emigration into the lungs. *IL17*-induced influx and reduced apoptosis of neutrophils result in their persistence in the lower airways of asthmatic horses (*100*). Neutrophil apoptosis is sometimes accompanied by the formation of NETs. While the release of antimicrobial peptides and various proteases through NETosis helps eliminate pathogens, it can also trigger tissue damage and sustain chronic inflammation (*101*). The dysregulation of genes associated with NETosis in the neutrophils of asthmatic horses agrees with the previous observations of excessive NETosis in the lungs of severely asthmatic horses (*102*). The increased NETosis function suggests a pro-inflammatory role of neutrophils in asthmatic horses, especially in the pro-inflammatory neutrophil subtype. Conversely, several DEGs also suggested an anti-inflammatory phenotype, particularly in the apoptotic neutrophil subtype. This could represent a protective mechanism aimed at preventing an excessive inflammatory response within the lungs. In summary, our findings confirm that BALF neutrophils from horses with SEA have a significant pro-inflammatory effect through increased neutrophil persistence and facilitated NET formation in the lungs. The concomitant anti-Th17 transcriptomic profile observed in apoptotic neutrophils indicates a parallel attempt to mitigate lung inflammation. This suggests neutrophils act as effectors rather than primary instigators of asthmatic lung inflammation. Targeting treatment specifically towards the pro-inflammatory neutrophil subtype could disrupt the self-perpetuating inflammatory circle while preserving the antimicrobial functions of the remaining neutrophil subtypes.

By employing scRNA-seq on BALF cells of horses with SEA, we were able to elucidate important underlying immune mechanisms of the disease. However, this study had certain limitations. One significant challenge when studying horses is the inadequate quality of the current reference annotation, necessitating manual annotation of the cell clusters, particularly for poorly defined cell subtypes. Nonetheless, the detection of previously identified cell types and subtypes in equine BALF (*13*, *21*) supports the reproducibility of cluster annotation. Some clusters, such as the “undetermined AMs” cluster, could not be confidently annotated. Further scRNA-seq studies and complementary techniques are required to provide conclusive insights.

ScRNA-seq is a relatively new technology that comes with computational challenges. One such challenge is the ability to detect and filter technical multiplets without removing biologically significant signals representing cell-cell complexes or new cell types with a dual lineage signature. In this study, we hypothesized that cluster Mo/Ma5 represented *bona fide* monocyte-lymphocyte complexes, supported by the presence of a similar transcriptomic signature in equine BALF cells (*13*, *21*) and human PBMCs (*35*, *36*). Although the existence of cellular complexes was confirmed in human PBMCs using imaging flow cytometry (*35*, *36*), validation in horses has yet to be performed. Another potential limitation associated with the 10X Genomics droplet-based technique is its low sensitivity for genes with a low average expression, which could explain the discrepancies with previous bulk RNA or proteomics studies, such as the absence of upregulated Th1 and Th2-associated cytokines.

This study was conducted on a small population. A power analysis indicated the need for a minimum of six control and six asthmatic horses with 500 cells/sample to achieve adequate statistical power. Unfortunately, the scRNA-seq experiment failed for one of the control samples. However, considering that we obtained more than 5,000 cells per sample, we are confident that the number of cells analyzed was sufficient to detect gene expression differences between the two groups.

Since this study specifically focused on neutrophilic SEA, our results cannot be generalized to other asthma subtypes. This is exemplified by a previous scRNA-seq study on equine BALF cells from horses with mastocytic asthma (*21*), which exhibited a different set of differentially regulated genes compared to our study. For example, *FKBP5* was significantly upregulated in the mast cells of asthmatic horses, while we did not detect this gene in our dataset. Hence, studying different endotypes separately is crucial to obtain meaningful results.

In conclusion, this scRNA-seq analysis of equine bronchoalveolar cells provided insights into the major immune mechanisms underlying severe equine asthma. The use of scRNA-seq allowed us to overcome the influence of varying cell type distribution associated with the disease and gain unprecedented resolution into the pathophysiology of SEA. This represents a significant breakthrough, challenging the prevailing perception of SEA as a Th2-associated disease. Consequently, the use of IgE-based tests and hyposensitization therapy in SEA should be reconsidered. We identified the crucial role of monocytes in initiating the Th17 response in the lungs, and the upregulation of *CXCL13* in lung and blood monocytes suggests its potential as a biomarker for SEA and as a therapeutic target. Our findings indicate that monocyte activation may occur through direct cell-cell contact, a hypothesis that should be tested using imaging flow cytometry. This has the potential to reshape our understanding of immunotherapy approaches. Notably, therapies targeting Th17-associated cytokines have proven ineffective in reducing symptoms in human asthma (*92*). One possible approach could be to prevent monocyte activation by targeting monocyte-T cell synapses. Our results demonstrate several parallels with previous studies on non-Th2 neutrophilic asthma in humans, further validating the horse as a valuable model for studying human asthma.

## MATERIALS AND METHODS

### Study design

In this observational case-control study, we recruited SEA-affected horses and controls based on their medical history. Following power analysis, we selected six asthmatic and six control horses, using BALF quality, respiratory symptom history and BALF neutrophilia as inclusion criteria. We applied 10X Genomics 3’-end scRNA-seq to the cryopreserved bronchoalveolar cells (∼ 6,000 cells/horse). The experiment failed in one control horse, leaving 11 horses for the data analysis. Our objectives were to assess the effect of SEA on i) the distribution of cell types and cell subtypes in the BALF and ii) the DGE within each of the cell types/subtypes identified.

### Study population

All animal experiments were performed according to the local regulations and with the consent of horse owners. This study was approved by the Animal Experimentation Committee of the Canton of Bern, Switzerland (BE4/20+). Privately owned horses were prospectively enrolled for a concomitant study on equine asthma (*104*). Suitable candidates were identified through a validated owner questionnaire (*105*, *106*). Requirements for study enrollment were an age ≥ 5 years old, a longer than 2-month history of being fed hay, no history of immunotherapy, no history of upper airway disease, no evidence of current systemic disease and a rectal temperature ≤ 38.5°C the day of the exam. Additionally, the horses should not have received any corticosteroids, bronchodilators or anti-histaminic administration nor suffered from a respiratory infection in the two weeks preceding the examination. Owners were asked to bring a healthy horse (without respiratory symptoms) from the same barn along with their asthmatic horse to the Bern ISME equine clinic. Six additional healthy horses belonging to the clinic employees or to an affiliated clinic (ISME Avenches) were enrolled. In total, 94 horses (46 healthy and 48 asthmatic) were examined. Horses underwent the following standard diagnostic procedures to characterize their respiratory status: physical examination, rebreathing examination, arterial blood gas analysis, lower airway endoscopy, tracheal aspirate and bronchoalveolar lavage. A BALF aliquot from 35 horses from the Warmblood breed and aged ≥ 6 years old was processed and stored for putative scRNA-seq.

### Power calculation

Data simulation was performed on a publicly available dataset to estimate the sample size required to attain sufficient statistical power. The dataset was originally published in a study aiming to characterize the immune cells in the peripheral blood of healthy horses using scRNA-seq (*107*) . We used scRNA-seq template data constructed based on the monocyte and dendritic cells from three healthy Warmblood horses (GSE148416). Data simulation was performed with Hierarchicell (*108*), which captures key characteristics from real datasets. Hierarchicell uses normalized data to obtain estimates of within sample variance (intra-individual variation) and between sample variance (inter-individual variation and dropout rates) by pruning genes to a set of uncorrelated genes. Using the estimated parameters from the template dataset, we simulated an RNA-seq expression matrix for 1,000 genes. The amount of fold change in expression between case and control groups was set at two. The number of cells per individual was set to be in the [150 – 1,000] range. The number of cases and controls were set to be in the [3 – 6] range. On the tSNE plot representing 6 controls and 6 cases with 500 cells/sample (see Suppl. fig. 5), the cases and controls are clearly separated. We thus elected to sequence 6 control samples and 6 SEA samples to reach adequate statistical power while optimizing sequencing cost.

### Case selection for scRNA-seq

Selection of the samples for scRNA-seq was made based on the BALF quality (yield ≥ 30% and foam indicating presence of pulmonary surfactant) and on the horses’ historical, clinical and laboratory features, with the goal of selecting the most archetypal phenotypes for both control and SEA groups. Horses included in the SEA group had a Horse Owner Assessed Respiratory Signs (HOARSI) score ≥ 3 (*105*, *109*) and a BALF neutrophilia > 10%. The control group consisted of horses with a HOARSI = 1 and a normal BALF differential cell count (< 10% BALF neutrophils, < 2% mast cells and < 1 % eosinophils).

### Sample collection

Details about horse examination, respiratory work-up and BALF collection have been published elsewhere (*104*).

### Cytology

Preparation of the slides for cytological analysis are described in a previous publication (*104*). The manual differential cell count including macrophages, lymphocytes, neutrophils, eosinophils and mast cells was performed on a minimum of 400 cells and on four different microscopic fields.

### Cryopreservation

The protocol used to freeze and subsequently thaw the BALF cells was adapted from the protocol we develop in our proof-of-concept study (*13*). The detailed laboratory protocol can be found in the supplementary materials. All samples were processed in less than two hours. Characteristics of the cell suspensions subject to scRNA-seq are provided in the supplementary table 14. For each sample, an aliquot of the final cell suspension was saved for the cytocentrifuge preparations.

### Single-cell cDNA library preparation and scRNA-seq

GEM generation & barcoding, reverse transcription, cDNA amplification and 3’ gene expression library generation steps were all performed according to the Chromium Next GEM Single Cell 3ʹ Reagent Kits v3.1 (Dual Index) User Guide (10x Genomics CG000315 Rev B) with all stipulated 10x Genomics reagents. Specifically, 12.0 µL of each cell suspension (1,100 cells/µL) and 31.2 µL of nuclease-free water were used for a targeted cell recovery of 8,000 cells. GEM generation and barcoding was followed by a GEM-reverse transcription incubation, a clean-up step and 1**1** cycles of cDNA amplification. The resulting cDNA was evaluated for quantity and quality using a Thermo Fisher Scientific Qubit 4.0 fluorometer with the Qubit dsDNA HS Assay Kit (Thermo Fisher Scientific, Q32851) and an Advanced Analytical Fragment Analyzer System using a Fragment Analyzer NGS Fragment Kit (Agilent, DNF-473), respectively. Thereafter, 3ʹ gene expression libraries were constructed using a sample index PCR step of 12-14 cycles. The generated cDNA libraries were tested for quantity and quality using fluorometry and capillary electrophoresis as described above. The cDNA libraries were pooled and sequenced with a loading concentration of 300 pM, asymmetric paired-end and dual indexed, on a shared **I**llumina NovaSeq 6000 sequencer using a NovaSeq 6000 S4 Reagent Kits v1.5 (200 cycles; **I**llumina, 20028313). The read set-up was as follows: read 1: 28 cycles, i7 index: 10 cycles, i5: 10 cycles and read 2: 90 cycles. The quality of the sequencing runs was assessed using **I**llumina Sequencing Analysis Viewer (**I**llumina version 2.4.7) and all base call files were demultiplexed and converted into FASTQ files using **I**llumina bcl2fastq conversion software v2.20. More than 50,000 reads/cell were generated for each sample. All steps were performed at the Next Generation Sequencing Platform, University of Bern.

### Data pre-processing

The raw fastq sequencing data was processed using the Cell Ranger (v6.0) standard workflow to generate a count matrix of gene expression values. The annotations for the 3’-untranslated regions of the genes in the reference genome (Equus caballus NCBI annotation release 103) were extended by 2 kb, following the methodology described in a previous study (*13*). Supplementary table 1 contains the summary metrics of the identified cells for the 12 samples.

### Quality control, doublet filtering and data normalization

Quality control was carried out with the R package Scater (v1.28.0) (*110*). Downstream analysis was conducted using the R package Seurat (v4.3) (*111*). One control sample did not meet the initial quality control criteria (only 844 cells retrieved), resulting in its exclusion. The output of CellRanger for the remaining 11 samples consisted of 75,727 cells. Doublet detection was conducted with scDblFinder (*112*), which led to the removal of 10,234 cells. Based on visual inspection, we further filtered 15,465 cells containing less than 200 genes or more than 8,000 gene features and/or greater than 15% mitochondrial genes. Overall, 25,699 (33.9%) cells were filtered prior to integration.

### Integration and data normalization

Data were integrated based on disease status (asthmatic vs control) using 3,000 integration features. Data normalization was conducted with the sctransform R package (v0.3.5) (*113*, *114*) using 3,000 variable features. The final dataset for downstream analysis contained 60,262 cells with 56,595 gene features.

### Principal component analysis (PCA) and cell clustering

Dimensionality reduction was performed using Principal Component Analysis (PCA). The optimal number of principal components (PCs) was based on an elbow plot. We conducted clustering on the first 16 PCs using the default Louvain algorithm (“FindNeighbors()” function in Seurat). Cluster granularity was explored using the clustree R package (*115*) and Uniform Manifold Approximation and Projection method (UMAP) in order to select the best clustering resolution. The Seurat function “FindCluster()” was run with a resolution of 0.6. Each of the six major cell populations were isolated using the “subset()” Seurat function. The previous steps were repeated to independently analyze B cells (11 PCs, clustering resolution 0.3), dendritic cells (DCs) (10 PCs, clustering resolution 0.1), mast cells (7 PCs, clustering resolution 0.4), monocyte-macrophages (Mo/Ma) (18 PCs, clustering resolution 0.2), neutrophils (11 PCs, clustering resolution 0.15) and T cells (18 PCs, clustering resolution 0.3).

### Cell cycle analysis

To analyze the effect of the cell cycle stage on clustering, we converted the human markers for the G2M and S phases (“cc.genes.updated.2019”) to their equine orthologs using the Biomart R package. Cells were divided into cycling (G2M phase) or resting (S phase) based on the score attributed by the “CellCycleScoring()” Seurat function.

### Automated cluster annotation

To facilitate cell cluster annotation, we performed automated annotation using the scSorter package (*116*). The annotation file was constructed with the top ten DEGs (output of “FindAllMarkers()” function) for each cluster identified in our previous equine BALF cells scRNA-seq analysis (*13*). Automated annotation was performed on the complete dataset and on the major cell groups for which previous annotation was available (B cells, Mo/Ma and T cells).

### Manual cluster annotation

For the complete dataset, the annotation of the major cell groups was confirmed by visualization of the gene expression pattern of canonical markers for cell type using the “DotPlot()”, “VlnPlot()” and “FeaturePlot()” functions in Seurat. Expression scores for cell-specific group of genes were calculated using the Seurat function “AddModuleScore()” to facilitate cell type identification. Annotation was further ascertained by investigation of the markers list provided by the “FindAllMarkers()” function, using an adjusted *P-*value <0.05 and an average log2 fold change > 0.25. The cellular specificity of the markers was inferred based on the Human Protein Atlas version 22.0 database (*117*). The subclusters within the major cell groups were manually annotated, as automated annotation did not allow for confident annotation.

### Differential gene expression (DGE) analysis

Differential gene expression analysis between the SEA and the control groups was performed with the R package Nebula (*118*). Nebula incorporates a negative binomial mixed model to consider the hierarchical structure of the data, decomposing the total overdispersion into between-subject and within-subject components. Only DEGs with an adjusted *P-*value <0.05 and an absolute average log2 fold change >1 were considered. The biological significance of the DEGs was scrutinized using the Human Protein Atlas database 22.0 database (*117*) and literature search (PubMed).

### Comparison of cellular distribution between groups

The cell type proportions were compared between control and asthma groups using the “propeller()” function from the speckle package in R (*119*). Propeller performs a variance stabilizing transformation on the matrix of proportions and fits a linear model for each cluster before implementing a moderated t-test to compare the groups. Benjamini and Hochberg false discovery rates are calculated to account for multiple testing of cell clusters. The level of significance was set at 0.05.

### Comparison of cellular distribution between sample types and counting techniques

Based on a variance analysis, the proportions of the five cytologically distinguishable leukocytes in the sample could be considered normally distributed, enabling parametric testing. To compare the leukocyte proportions, we conducted a repeated measure ANOVA, considering individual horses and the analysis type (BALF cytology, cell suspension cytology or cell suspension scRNA-seq) as predictor variables. Specifically, we compared the leukocyte proportions between i) BALF cytology and cell suspension cytology, and ii) cell suspension cytology and scRNA-seq. The level of significance was set at 0.05.

## Acknowledgments

The authors would like to thank Dr. med. vet. Michelle Wyler and Dr. med. vet. Nicole Altermatt for their assistance in sample collection. We thank Dr. Pamela Nicholson and the laboratory technicians from the Next Generation Sequencing Platform of the University of Bern for performing the sequencing experiments. We are grateful for the Interfaculty Bioinformatics Unit of the University of Bern for providing the high-performance computing infrastructure. The authors furthermore acknowledge Dr. med. vet. Laureen Peters for conducting cytological analysis and the laboratory technicians of the Central Diagnostic Laboratory of the Vetsuisse Faculty for their assistance in sample processing.

## Author contributions

SS, TL and VG designed the study. Funding was acquired by VG. SS collected and processed the samples. PN supervised the scRNA-seq experiments. SS and VJ performed the computational analysis. SS wrote the original draft. All authors reviewed, edited and approved the final version of the manuscript.

## Data and materials availability

The datasets generated for this study will be deposited in the European Nucleotide Archive (ENA) repository https://www.ebi.ac.uk, after publication in a peer-reviewed journal. The R code used for data analysis will be published.

## SUPPLEMENTARY MATERIALS

### Supplementary figures

**Supplementary figure 1.** Ribosomal protein genes differential expression across the seven T cell populations (n=30,005 cells) visualized with UMAP (A) and with a violin plot (B).

**Supplementary figure 2.** RNA feature count (A) and mitochondrial read percentage (B) across the six monocyte-macrophage (Mo/Ma) cell populations (n=22,370 cells).

**Supplementary figure 3.** Ribosomal protein genes differential expression across the six monocyte-macrophage (Mo/Ma) cell populations (n=22,370 cells) visualized with UMAP (A) and with a violin plot (B).

**Supplementary figure 4.** Distributions of the five cytologically distinguishable leukocytes obtained with cytology on bronchoalveolar lavage fluid (BALF), with cytology on the cell suspension (post cryopreservation) and with scRNA-seq on the cell suspension. T cells and B cells are counted as lymphocytes, while Mo/Ma and DCs are counted as macrophages.

**Supplementary figure 5.** tSNE visualization of the data simulation using 6 control and 6 cases with 500 cells per sample. Hierarchicell [PMID: 33932993] was used to simulate the data based on scRNA-seq template data from healthy Warmblood horses (GSE148416). A 1000-gene RNA-seq matrix was generated with a fold change of 2 to simulate differential expression between case and control groups. Varying the number of cells (150 to 1,000) and samples (3 to 6) showed distinct clustering when using 6 horses per group with 500 cells per sample.

### Supplementary tables

**Supplementary table 1.** Summary metrics of the detected cells for the 12 samples sequenced **Supplementary table 2.** Marker genes for the 19 clusters (complete dataset)

**Supplementary table 3.** Marker genes for the six major cell types (complete dataset)

**Supplementary table 4**. Marker genes for the neutrophil clusters (independent reanalysis)

**Supplementary table 5.** Marker genes for the B cell clusters (independent reanalysis)

**Supplementary table 6.** Marker genes for the T cell clusters (independent reanalysis)

**Supplementary table 7.** Marker genes for the monocyte-macrophage clusters (independent reanalysis)

**Supplementary table 8.** Marker genes for the dendritic cell clusters (independent reanalysis)

**Supplementary table 9.** Differential gene expression analysis – Major cell types

Supplementary table 10. Differential gene expression analysis – Neutrophils

Supplementary table 11. Differential gene expression analysis – T cells

Supplementary table 12. Differential gene expression analysis – Monocytes-Macrophages

Supplementary table 13. Differential gene expression analysis – Dendritic cells

Supplementary table 14. Characteristics of the cell suspensions subject to scRNA-seq

### Supplementary material and methods

Experimental protocol for equine bronchoalveolar cells cryopreservation

