## Supplementary material for "Single-cell profiling of bronchoalveolar cells reveals a Th17 signature in neutrophilic severe equine asthma": Supplementary figures.pdf

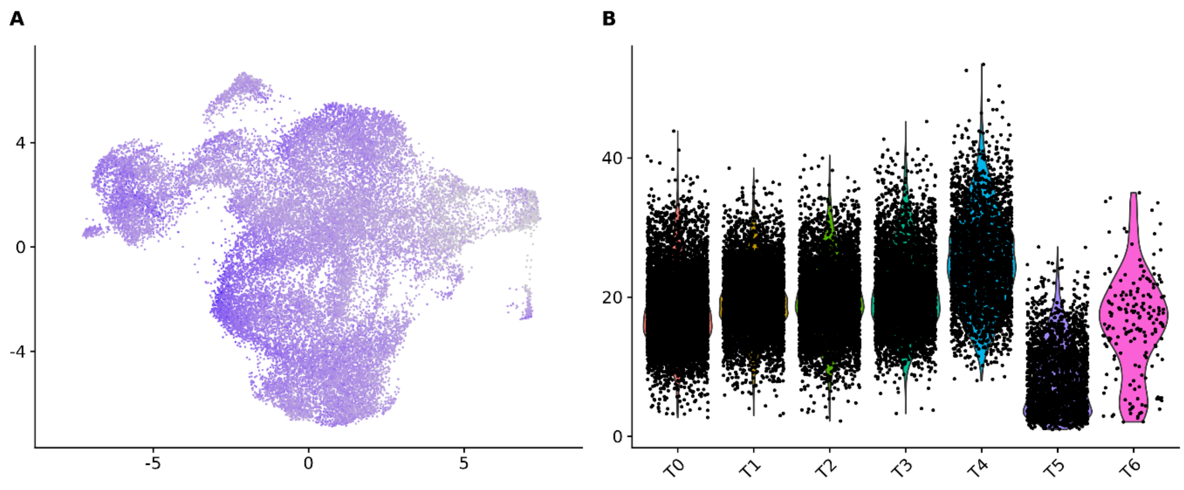


**Supplementary figure 1.** Ribosomal protein genes differential expression across the seven T cell populations (n=30,005 cells) visualized with UMAP **(A)** and with a violin plot **(B).**

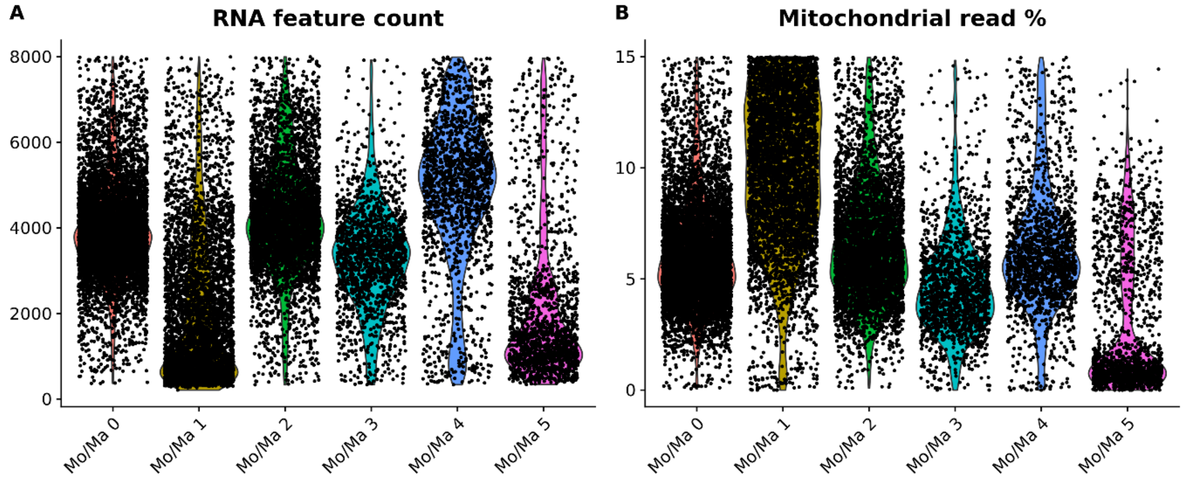


**Supplementary figure 2.** RNA feature count **(A)** and mitochondrial read percentage **(B)** across the six monocyte-macrophage (Mo/Ma) cell populations (n=22,370 cells).

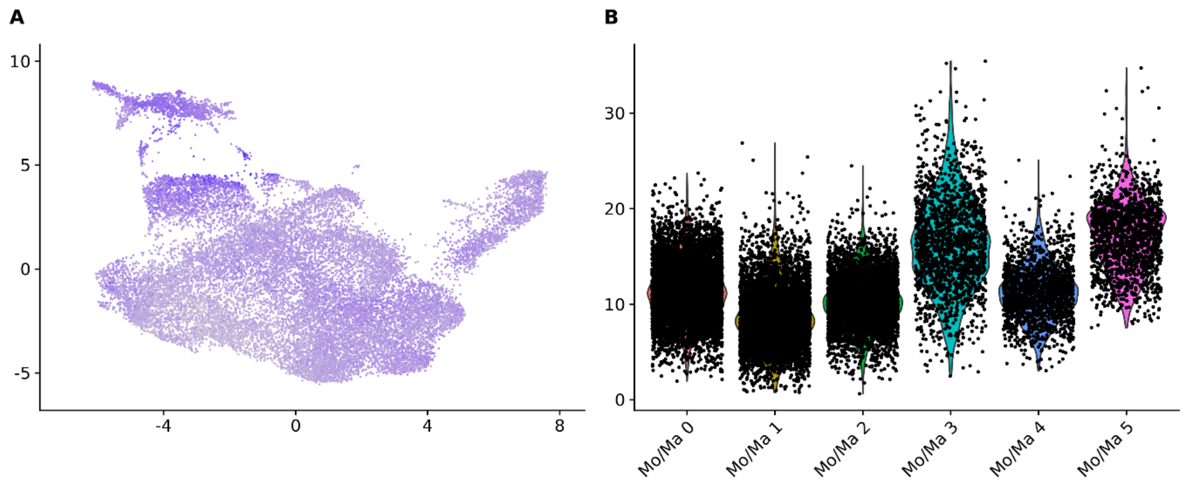


**Supplementary figure 3.** Ribosomal protein genes differential expression across the six monocyte-macrophage (Mo/Ma) cell populations (n=22,370 cells) visualized with UMAP **(A)** and with a violin plot **(B)**.

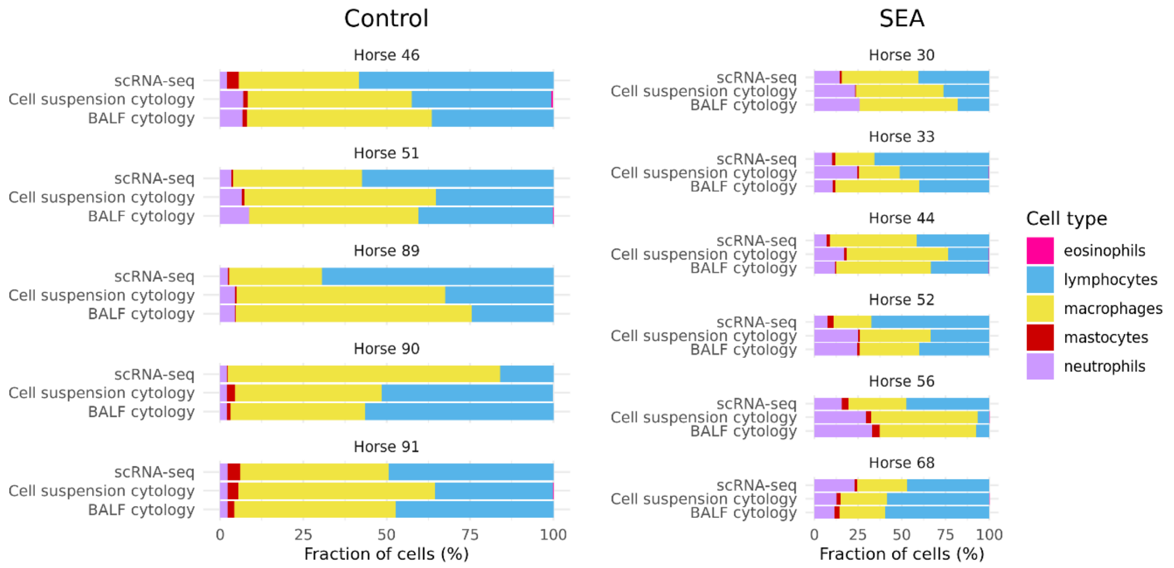


**Supplementary figure 4.** Distributions of the five cytologically distinguishable leukocytes obtained with cytology on bronchoalveolar lavage fluid (BALF), with cytology on the cell suspension (post cryopreservation) and with scRNA-seq on the cell suspension. T cells and B cells are counted as lymphocytes, while Mo/Ma and DCs are counted as macrophages.

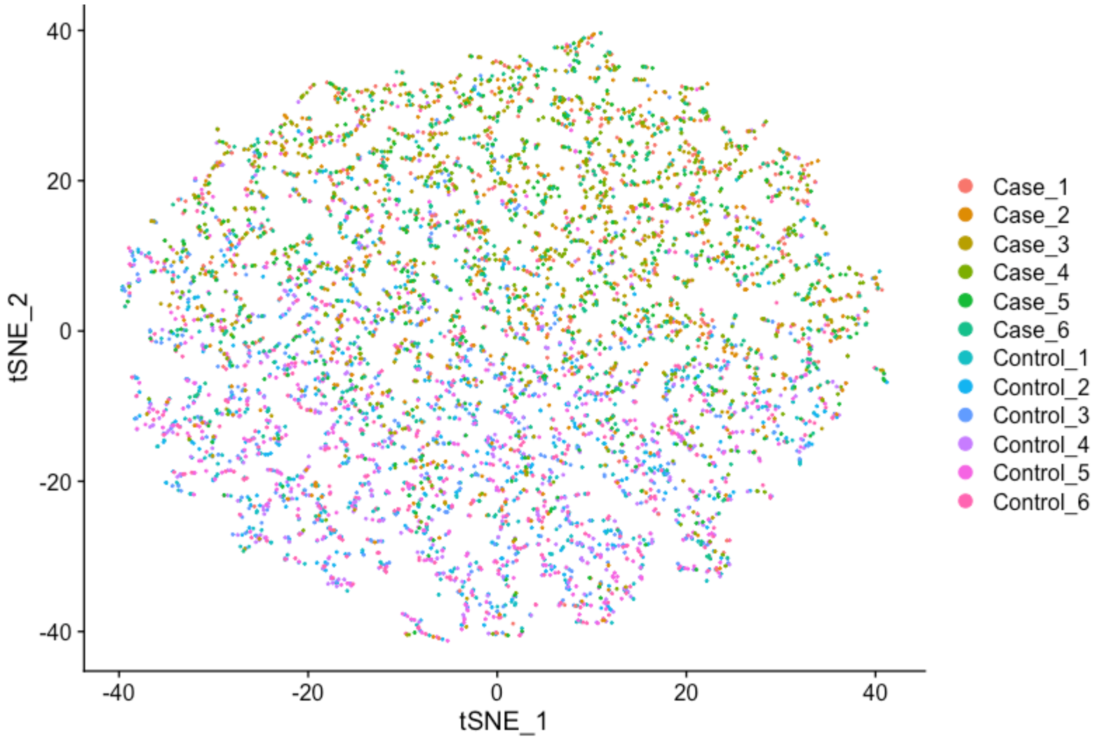


**Supplementary figure 5.** tSNE visualization of the data simulation using 6 control and 6 cases with 500 cells per sample. Hierarchicell [PMID: 33932993] was used to simulate the data based on scRNA-seq template data from healthy Warmblood horses (GSE148416). A 1000-gene RNA-seq matrix was generated with a fold change of 2 to simulate differential expression between case and control groups. Varying the number of cells (150 to 1,000) and samples (3 to 6) showed distinct clustering when using 6 horses per group with 500 cells per sample.
