## Supplementary material for "Single-cell profiling of bronchoalveolar cells reveals a Th17 signature in neutrophilic severe equine asthma": Supplementary material and methods - Experimental protocol for equine bronchoalveolar cells cryopreservation .pdf

### *EXPERIMENTAL PROTOCOL: cryopreservation of equine bronchoalveolar cells for single-cell mRNA sequencing*

#### 1 Overview

This protocol outlines cryopreservation and thawing of equine bronchoalveolar (BAL) cells for use with 10x Genomics (Pleasanton, CA, USA) Single Cell protocols. This experimental protocol is adapted from the protocol CG00039 Rev D protocol validated by 10X Genomics for the cryopreservation of human peripheral blood mononuclear cells. Minor changes were made to the protocol provided in the supplementary material of Sage et al. 2021.

#### 2 Sample type

This protocol is suitable for fresh equine BAL cells. Freshly collected BAL fluid should be filtered through a 40-µm cell strainer (BD Falcon™, Biosciences, USA) and kept on ice until processing.

#### 3 Reagents & Consumables

| Vendor | Item | Part number |
|---|---|---|
| Fisher Scientific | Corning™ CoolCell™ 5 mL LX Cell Freezing Container | FS 15592771<br>Corning™ 432005 |
| | Eppendorf™ DNA 5 mL LoBind Tubes | FS 16170371<br>Eppendorf™ 0030122348 |
| | Eppendorf™ Snap-Cap Microcentrifuge Flex-Tube™ 1.5 mL Tubes | FS 13094697<br>Eppendorf™ 02550-07 |
| | Falcon™ 15 mL Conical Centrifuge Tubes | FS 11507411<br>Falcon™ 352196 |
| | Falcon™ 50 mL Conical Centrifuge Tubes | FS 10788561<br>Falcon™ 352070 |
| | MP Biomedicals dimethylsulfoxid (DMSO) ≥99 % | FS 11440522<br>MPB 0219141880 |

| | | |
|---|---|---|
| OMNI Life Science | CASY Cell Counter & Analyzer | CASY TTT |
| | CASYton | |
| | CASYcups | |
| Sigma-Aldrich | Roche® Protector RNase Inhibitor 40 U/µL | SA 3335402001 |
| Thermo Fisher Scientific | Gibco™ Dulbecco's phosphate-buffered saline (DPBS), no calcium, no magnesium | TFS 14190094 |
| | Gibco™ Fetal Bovine Serum (FBS), certified | TFS 16000044 |
| | Gibco™ RPMI 1640 Medium | TFS 11875093 |
| | Invitrogen™ UltraPure™ BSA (50 mg/mL) | TFS AM2616 |
| | Nunc™ Biobanking and Cell Culture 4.5 mL Cryogenic Tubes | TFS 379146 |

#### 4 Media composition

| **Cryopreservation** | |
|---|---|
| **Media** | **Composition** |
| Resuspension Medium (maintain at 4°C) | 40% FBS in RPMI with RNAse inhibitor 1 U/µL |
| Freezing Medium (maintain at 4°C) | 30% DMSO in RPMI with 40% FBS |
| **Thawing & Resuspension** | |
| **Media** | **Composition** |
| Complete Growth Medium (maintain at 37°C) | 10% FBS in RPMI |
| Resuspension Solution (maintain at room temperature) | 0.04% BSA in DPBS with RNAse inhibitor 0.8 U/µL |

#### 5 Protocol

##### 5.1 Cryopreservation

Pre-cool a cell freezing container by placing it on ice before starting cryopreservation.

a. Place cells on ice.

b. Gently mix the cells.

c. Determine cell viability and total cell number. If using the CASY counter & analyzer, add 50 µL of the cell suspension to a CASYcup filled with 10 ml CASYton. Invert the cup slowly 3 times.

d. Centrifuge at 300 rcf for 5 min at 4°C.

e. Remove the supernatant.

f. Resuspend the cell pellet in an appropriate volume of chilled Resuspension Medium to achieve a concentration of 20 x $10^6$ cells/mL. Maintain the cells on ice.

g. Add an equivalent volume of chilled Freezing Medium to achieve a concentration of 10 x $10^6$ cells/mL. Gently mix the cells.

h. Dispense cell suspension aliquots into the pre-cooled cryovials and place the cryovials inside a pre-cooled cell freezing container.

i. Place the cell freezing container in a −80°C freezer for ≥ 4 h. After 4 h, transfer the cryovials to a cryobox in the −80°C freezer.

##### 5.2 Thawing & Resuspension

Set up a water bath to 37°C before starting cell thawing. All cell washes are performed at room temperature.

a. Remove cryovials from storage and immediately thaw in the water bath at 37°C for 2-3 min. Do not submerge the entire vial in the water bath. Remove from the water bath when a tiny ice crystal remains.

b. In a biosafety hood, slowly transfer thawed cells to a 50-mL conical tube using a wide-bore pipette tip. Rinse the cryovial with 1 mL warm Complete Growth Medium and add the rinse dropwise (1 drop per 5 sec) to the 50-mL conical tube while gently shaking the tube.

c. Sequentially dilute cells in the 50-mL conical tube by incremental 1:1 volume additions of medium for a total of 5 times (including dilution at step b). Add medium at a speed of 1 mL/3-5 sec to the tube and swirl.

d. Centrifuge at 300 rcf for 5 min at 20°C.

e. Remove most of the supernatant, leaving ~1 mL and resuspend cell pellet in this volume using a regular-bore pipette tip.

f. Add an additional 9 mL Complete Growth Medium (at a speed of 1 mL/ 3-5 sec) to achieve a total volume of ~10 mL.

g. Determine the cell concentration.

h. Transfer 10 x $10^5$ cells into a new 15-mL tube.

i. Centrifuge at 300 rcf for 5 min at 20°C.

j. Remove the supernatant without disrupting the cell pellet.

k. Using a wide-bore pipette tip, add 200 µL of Resuspension Solution and gently pipette mix 5 times.

l. Transfer the cells into a 1.5-mL DNA LoBind microcentrifuge tube. Rinse the 15-mL conical tube with 200 µL Resuspension Solution and transfer the rinse into the 1.5-mL DNA LoBind tube containing cells. Gently pipette mix 1 time.

m. Centrifuge at 300 rcf for 5 min at 20°C.

n. Remove the supernatant without disrupting the cell pellet.

o. Add an appropriate volume of Resuspension Solution to obtain a concentration of 0.7-1.2 x $10^6$ cells/mL (approx. 300 µL). Gently pipette mix using a regular-bore pipette tip until a single cell suspension is achieved.

p. Determine cell concentration and viability. The targeted final concentration is 0.7-1.2 x $10^6$ cells/mL.

q. Once the final cell concentration is achieved, place cells on ice.

r. Proceed immediately to the 10x Genomics Single Cell protocol.

*s.* Keep the remaining cell suspension on ice until processing for cytology.
