## Supplementary material for "Single-cell profiling of bronchoalveolar cells reveals a Th17 signature in neutrophilic severe equine asthma": Supplementary tables.pdf

**Supplementary table 1.** Summary metrics of the detected cells for the 12 samples sequenced

| | **A30** | **A33** | **A44** | **C46** | **C51** | **A52** | **A56** | **A68** | **C87** | **C89** | **C90** | **C91** |
|---|---|---|---|---|---|---|---|---|---|---|---|---|
| Estimated number of cells | 5,102 | 5,609 | 7,201 | 6,205 | 7,910 | 7,324 | 7,842 | 8,085 | 844 | 6,915 | 6,550 | 6,112 |
| Fraction reads in cells (%) | 91.7 | 89.2 | 91.0 | 87.6 | 93.7 | 90.6 | 93.7 | 89.2 | 34.6 | 90.7 | 76.6 | 95.1 |
| Mean reads per cell | 106,308 | 98,268 | 76,638 | 99,849 | 78,917 | 96,844 | 77,905 | 95,081 | 627,770 | 96,555 | 93,097 | 84,021 |
| Median UMIs per cell | 2,306 | 1,801 | 3,940 | 1,471 | 1,912 | 1,747 | 1,955 | 2,087 | 1,581 | 1,199 | 1,662 | 2,511 |
| Median genes per cell | 896 | 777 | 1,351 | 664 | 778 | 734 | 839 | 814 | 778 | 589 | 699 | 1,042 |
| Total genes detected | 14,464 | 16,512 | 17,109 | 16,513 | 17,018 | 17,429 | 17,050 | 17,113 | 14,442 | 16,130 | 16,249 | 17,111 |
| Sequencing saturation (%) | 73.8 | 78.9 | 59.6 | 77.2 | 63.7 | 78.7 | 68.8 | 74.7 | 86.7 | 83.2 | 77.4 | 60.3 |
| Reads mapped confidently to genome (%) | 90.2 | 91.3 | 91.6 | 90.6 | 90.3 | 90.8 | 91.0 | 89.4 | 88.0 | 89.1 | 90.1 | 91.0 |
| Reads mapped confidently to exonic regions (%) | 48.8 | 44.9 | 41.1 | 42.0 | 40.1 | 40.2 | 40.2 | 39.4 | 33.6 | 46.2 | 40.8 | 41.8 |
| Reads mapped confidently to transcriptome (%) | 44.4 | 40.8 | 37.1 | 37.7 | 36.2 | 36.2 | 36.1 | 35.4 | 29.9 | 41.7 | 36.3 | 37.7 |

*Data generated with CellRanger v6.0 using EqCab 3.0 NCBI annotation release 103 with 3'-UTR regions extended by 2 kb. A: asthmatic horse; C: control horse*
